# Genomic data provides new insights on the demographic history and the extent of recent material transfers in Norway spruce

**DOI:** 10.1101/402016

**Authors:** Jun Chen, Lili li, Pascal Milesi, Gunnar Jansson, Mats Berlin, Bo Karlsson, Jelena Aleksic, Giovanni G. Vendramin, Martin Lascoux

## Abstract

Primeval forests are today exceedingly rare in Europe and transfer of forest reproductive material for afforestation and improvement have been very common, especially over the last two centuries. This can be a serious impediment when inferring past population movements in response to past climate changes such as the last glacial maximum (LGM), some 18,000 years ago. In the present study, we genotyped 1,672 individuals from three Picea species (P. abies, P. obovata, and P. omorika) at 400K SNPs using exome capture to infer the past demographic history of Norway spruce and estimate the amount of recent introduction used to establish the Norway spruce breeding program in Southern Sweden. Most of these trees belong to P. abies and originate from the base population of the Swedish breeding program. Others originate from populations across the natural ranges of the three species. Of the 1,499 individuals stemming from the breeding program, a large proportion corresponds to recent introductions. The split of P. omorika occurred 23 million years ago (mya), while the divergence between P. obovata and P. abies began 17.6 mya. Demographic inferences retrieved the same main clusters within P. abies than previous studies, i.e. a vast northern domain ranging from Norway to central Russia, where the species is progressively replaced by Siberian spruce (P. obovata) and two smaller domains, an Alpine domain, and a Carpathian one, but also revealed further subdivision and gene flow among clusters. The three main domains divergence was ancient (15 mya) and all three went through a bottleneck corresponding to the LGM. Approximately 17% of P. abies Nordic domain migrated from P. obovata ~103K years ago, when both species had much larger effective population sizes. Our analysis of genome-wide polymorphism data thus revealed the complex demographic history of Picea genus in Western Europe and highlighted the importance of material transfer in Swedish breeding program.

## Introduction

Plant and animal species in Western Europe were exposed repeatedly to ice ages and therefore went through cycles of contraction and expansion from one or multiple refugia (Petit *et al.* 2003; Depraz *et al.* 2008; Schmitt and Haubrich 2008; Pilot *et al.* 2014). The dominant paradigm of early phylogeographic studies postulated that species survived the last glacial maximum (LGM) in small refugia in Southern Europe (Taberlet *et al.* 1998; Hewitt 2000; Tzedakis *et al.* 2002; Tzedakis *et al.* 2003) and then recolonized Europe through different routes. However, while this still seems to be true for temperate species such as oaks (Petit *et al.* 2003), paleo-ecological data (pollen fossils maps, macrofossils) as well as genetic studies indicated that boreal species were able to survive at much higher latitudes than initially foreseen. This generally led to a more diffuse and complex population genetic structure than in species with southern refugia (Willis *et al.* 2000; Lascoux *et al.* 2004; Tzedakis *et al.* 2013). In the case of genetic studies, care was generally taken to sample in natural forests, though in some cases this turned out to be difficult. For instance, in larch *(Larix decidua)* populations in central Europe, some of the discordant phylogeographic patterns corresponded to recent translocations by German immigrants of seeds from stands belonging to a different refugium (Wagner et al. 2015).

Norway spruce *(Picea abies)* is a dominant conifer tree species in Western Europe whose current distribution is divided into three main areas: a vast northern domain ranging from Norway to central Russia, where the species is introgressed and progressively replaced by Siberian spruce *(P. obovata)* and two smaller domains, an Alpine domain, and a Carpathian one (Fig. 1; EUFORGEN 2009). Earlier population genetic and phylogeographic studies based on isozymes and cytoplasmic markers, respectively, suggested that this current distribution originated from two main LGM refugia: one centered in Russia, and another in the Alps (Tollefsrud *et al.* 2008; Tollefsrud *et al.* 2009). However, more recent studies based on nuclear sequence data suggested that northern populations were extensively admixed, unlike the southern ones that were rather homogeneous (Chen *et al.* 2012a; Tsuda *et al.* 2016). In particular, introgression from Siberian spruce *(P. obovata)* into Norway spruce could be detected as far as Central Russia, especially in the north, resulting in a large hybrid zone centered on the Urals.

**Figure 1.**
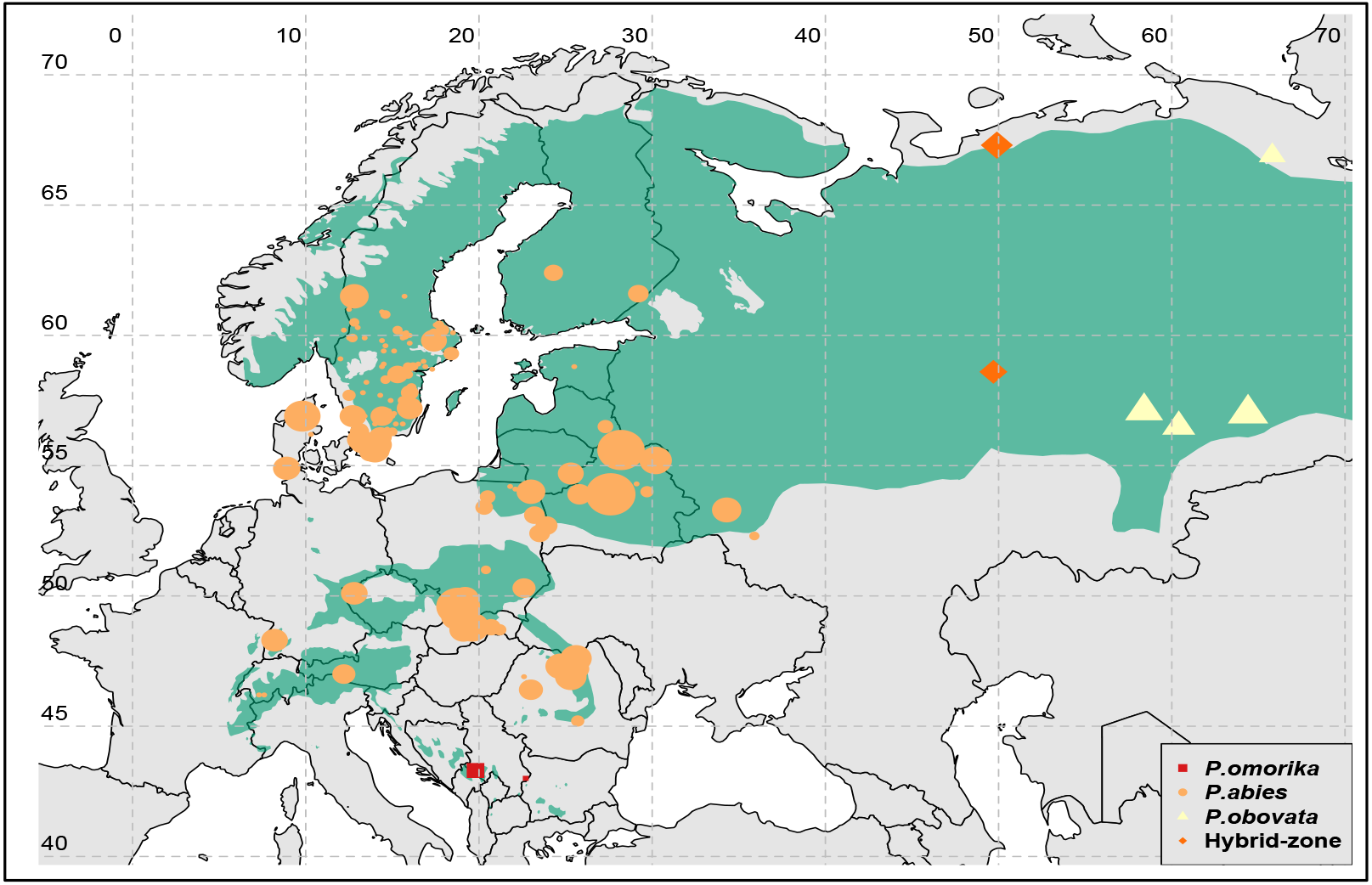
Distribution range of the three European spruce species (modified from EUFORGEN 2009, www.euforgen.org). Points are sampling coordinates of *P. obies* (orange discs), *P. obovata* (yellow triangles), *P. omorika* (red squares), the hybrid populations (gold diamonds); point sizes are proportional to the number of individuals in each location. For *P. obies*, they consist of both natural population samples and mother trees from common garden trials for the Swedish breeding program.

Furthermore, there is an apparent conflict in the relationship between northern populations of Norway spruce, southern ones and Siberian spruce at nuclear (SSR) and mitochondrial markers. SSR markers define two well-differentiated groups: southern and northern Norway spruce, on the one hand, and Siberian spruce, on the other hand (Tsuda *et al.* 2016). Mitochondrial DNA, in contrast, singles out southern populations of Norway spruce and grouped northern and Siberian spruce populations together (Lockwood *et al.* 2013; Tsuda *et al.* 2016). The latter led Lockwood *et al.* (2013) to propose that southern populations of Norway spruce be regarded as a different species or, at least, as a subspecies. Based on the joint pattern at nuclear and mtDNA, Tsuda *et al.* (2016) suggested that the difference in pattern could be explained by asymmetric migration of pollen and seed. Their study also confirmed the extent of introgression from *P. obovata* into northern *P. abies* populations and linked it to the post-glacial recolonization process. Two migration barriers were revealed: one in the Western Urals between the northern domain of *P. abies* and *P. obovata* and a second one between the Alps and Carpathian Mountains (Tsuda *et al.* 2016). The presence of three *P. abies* domains has been confirmed by studies using variation in cone morphology (Borghetti *et al.* 1988), organelle DNA markers (Achere *et al.* 2005; Tollefsrud *et al.* 2008), AFLP (Achere *et al.* 2005), sequence data (Heuertz *et al.* 2006) and genome-wide restriction site DNA markers (Fagernäs 2017).

Although based on a limited number of nuclear DNA markers, previous studies also indicated extensive shared ancestral polymorphisms, suggesting a relatively recent divergence time measured on an effective population size timescale, as well as weak but significant effect of migration (Heuertz *et al.* 2006; Chen *et al.* 2010; Li *et al.* 2010a; Chen *et al.* 2016; Tsuda *et al.* 2016). Chen *et al.* (2016) concluded that the Fennoscandian domain split from the two southern domains of *P. abies* around 5 million years ago (mya), i.e. before the Pliocene-Quaternary glaciation, which is consistent with estimates of dating based on the molecular clock (~6 mya, (Lockwood *et al.* 2013)). However, previous studies were limited to fairly simple demographic scenarios, such as “Isolation-with-Migration” models and, in particular, could not distinguish post-speciation contact from migration. None also considered translocations, and samples were generally assumed to be of local origin. Based on historical records on seed used in reforestation, the problem of transfer of reproductive material should be particularly acute in populations of Norway spruce, especially in southern Sweden. During the twentieth century, Sweden imported more than 210 tons of seed and more than 600 million plants, primarily for afforestation in Southern Sweden. This material initially came primarily from central Europe but interest later shifted eastwards with introduction of material from Belarus, the Baltic states and Romania that proved to be more resistant to frost than central European provenances (Myking *et al.* 2016).

The release of the Norway spruce genome and the development of reduced genome re-sequencing technologies (e.g. RAD-Seq, exome capture) provide us with the opportunity to investigate genome-wide pattern of diversity in hundreds or even thousands of individuals. Using whole genome polymorphism data, model-based and non-parametric clustering methods, allow detecting and quantifying subtle genetic admixture and migration (see (Alexander *et al.* 2009; Pickrell and Pritchard 2012; Galinsky *et al.* 20Ma16), for examples in humans) while coalescent or diffusion-based methods that use the joint site frequency spectrum (SFS) across multiple populations allow testing different demographic scenarios (Gutenkunst *et al.* 2009; Excoffier *et al.* 2013).

In this paper we thus investigated past demographics and recent translocations in Norway spruce using genome-wide SNP data from > 1,600 individuals sampled from: i) populations from southern Sweden that were used to establish the Swedish Norway spruce breeding program, ii) natural populations of *P. abies* across its natural range and iii) two close relatives, the Siberian spruce (*P. obovata*) and the Serbian spruce *(P. omorika).*

## Material and Methods

### Sample collection

Samples in this study consist of 1,672 individuals of three Picea species *(P. abies, P. obovata*, and *P. omorika)* with origins extending from 43.0°N to 67.3°N in latitude and from 7.3°E to 65.8°E in longitude (Fig. 1). The samples were gathered through two main channels. First, we collected needles from 1,499 individuals that were initially used to create the Swedish breeding program, i.e. were selected as “plus trees” (trees of outstanding phenotype) in 20-40 years old production forestry stands or selected as “superior” 3-4 years old seedling genotypes in commercial nurseries. This was done across central and Southern Sweden. Of those, 575 individuals had no clear records of their geographical origin; after genotypic clustering analyses, fifteen of them showed high similarity to *P. omorika* and were thus treated as such in the following analyses. Second, seedlings from individuals coming from natural populations (six *P. omorika*, 53 *P. obovata*, 74 *P. abies*, and 40 hybrid individuals) were collected after seed-germination in growth chambers in 2015. Those individuals were used as reference for genetic cluster definition and genotype assignation. *P. obovata* samples were collected from four Siberian populations: three along a longitudinal gradient in Southern Urals: Shalya (57.14°N, 58.42°E), Ekaterinburg (56.50°N, 60.35°E) and Tugulym (57.03°N, 64.37°E); and one in the Northern Urals: Krasnij Kamenj (66.54°N, 65.45°E). Two additional populations that are part of the *P. abies-P. obovata* hybrid zone were also included: Indigo, from high latitude (67.27°N, 49.88°E) and Kirov, from a lower latitude (58.60°N, 49.68°E); the former having a more even contribution of the two parental species (Tsuda *et al.* 2016). In comparison to these two continental species, *P. omorika* has today a very tiny distribution range that is confined to mountain regions of Western Serbia and Eastern Bosnia and Herzegovina.

### SNP identification and estimation of genetic diversity

Genomic DNA was extracted either from needles or buds in the case of the individuals from the breeding program or from needles from seedlings in the case of individuals sampled from natural populations using the DNeasy Plant Mini Kit (QIAGEN, Germantown, MD). 40,018 probes of 20bp long were designed to cover 26,219 *P. abies* gene modules (see more in Vidalis *et al.* 2018). Library preparation and exome-capture sequencing were performed by RAPiD Genomics, U.S.A. Paired-end short reads were aligned to *P. abies* genome reference v1.0 (Nystedt *et al.* 2013) using BWA-mem with default parameters (Li and Durbin 2009). We extracted the uniquely aligned properly paired reads in the scaffolds/contigs, where probes were designed to bait for. PCR duplicates were removed with PICARD v1.141 (http://broadinstitute.github.io/picard) and INDEL realignment was performed by GATK (McKenna *et al.* 2010). HaplotypeCaller was used for individual genotype identification and joint SNP calling was performed across all samples using GenotypeGVCFs. We then applied the same variant quality score recalibration protocols as Baison *et al.* (2018), which were trained on a set of ~21,000 SNPs identified from a pedigree study (Bernhardsson *et al.* 2018). This resulted in 2,406,289 SNPs after recalibration. SNPs were filtered following two criteria: (i) each allele of individual genotype should be called with more than two reads and (ii) more than half of the individuals should be successfully genotyped at each site. Five individuals were removed from our data due to insufficient read coverage. In total, 1,004,742 SNPs were retained for further analyses. Of these, 364,034 fall in the exons of 25,569 transcripts, 135,548 are synonymous and 228,486 cause amino acid changes. Of the remaining sites, 448,698 SNPs are in introns and 192,010 fall in intergenic regions.

Genetic diversity was calculated at all sites and also at 0-fold (coding sites at which all changes are nonsynonymous) and 4-fold sites (coding sites at which all changes are synonymous, π_0_ and π_4_, respectively). Their ratio was then calculated for protein coding sequences of *P. omorika, P. obovata*, and the three main domains of *P. abies* (Fennoscandian, Alpine, and Carpathian). Tajima’s D (Tajima 1989) and pairwise population fixation indices *F_ST_* between species and the main domains of *P. abies* (see below) were also calculated using polymorphisms in noncoding regions. The above summary statistics were calculated with custom Perl scripts.

### Population genetic clustering

Haplotype phasing was conducted using MACH v 1.0 (Li *et al.* 2010b) with default parameter setting (average switch error rate 0.009), pairwise linkage disequilibrium (LD) was then calculated using the HaploXT program in the GOLD package (Abecasis and Cookson 2000). For population clustering and admixture inference, the analyses were applied on 399,801 noncoding SNPs with significantly linked sites removed (pairwise LD ≥ 0.2 and FDR value ≤ 0.05).

EIGENSOFT v6.1.4 (Galinsky *et al.* 2016) was used to perform PCA on the genetic variation of *P. abies* and *P. obovota. P. omorika* was excluded from this PCA analysis due to its extremely high divergence from the other two species. For unsupervised population clustering, ADMIXTURE v1.3 (Alexander *et al.* 2009) was used with five-fold cross validation and 200 bootstrap replicates. The number of ancestral clusters (K) varied from one to eight and the best value was chosen at the lowest value of crossvalidation error (Fig. S1).

### Geographic inference for individuals with unclear sources

Geographic origin of the 575 “Unknown” individuals for which no confident records of geographical origin were available was assessed based on their genotype similarity to ascertained individuals. *P. abies* individuals of known origin were first grouped into seven major clusters based on genetic clustering results and their origin. These individuals were used as the training dataset in a “Random Forest” regression model implemented in R software (Liaw Andy and Matthew 2002). The first five components of a PCA analysis were used for model fitting and to classify the “Unknown” individuals into each of the seven clusters. Five-fold cross-validations were performed for error estimation. “Unknown” individuals were then assigned to the various genetic clusters defined from individuals from known origin. The whole regression process was repeated 1,000 times in order to estimate the confidence of each assignment.

### Admixture inference

TreeMix v1.13 (Pickrell and Pritchard 2012) was used to infer the direction and proportion of admixture events. A maximum likelihood phylogenetic tree was first built for the seven *P. abies* population clusters, *P. obovata*, and their hybrids by bootstrapping over blocks of 100 SNPs. *P. omorika* was used as an outgroup to root the tree. Admixture was tested between each pair of populations and branches were rearranged after each significant admixture event added to the tree. The number of admixture events was estimated by minimizing the residual matrix of model compared to observed data. To avoid over-fitting, we stopped adding admixture when the tree model explained over 95% of the variance.

### Demographic inference using multidimensional site frequency spectra (SFS)

Effective population sizes and divergence times between the three most divergent *P. abies* clusters (i.e., the Alpine, Carpathian and Fennoscandian) were first estimated. *P. omorika* was used to polarize the derive allele frequency and shared polymorphic sites between *P. omorika* and target populations were excluded for stringency. In pilot runs to maximize the likelihood for individual SFS, we noted that estimates of historical *N_e_* for all three populations are much larger than current ones suggesting that current populations had gone through bottlenecks. Therefore, a divergence model was applied with all three populations going through a sudden size contraction at *T*_Bot_. The Fennoscandian domain first split at time *T*_FAC_, followed by the split of the Alpine and Carpathian domains at *T*_AC_. To reduce model complexity, we used a constant migration rate (m = 1×10^-6^) between all pairs of populations averaged along the whole divergence history. To examine if migration could improve model composite likelihood, models with m = 0 and m=1×10^-6^ were compared based on Akaike’s weight of evidence. The aforementioned parameters were estimated by maximizing composite likelihoods based on observed 3-dimensional joint SFS using Fastsimcoal2 v2.6.0.2 (Excoffier *et al.* 2013). We performed 100 iterations of parametric bootstrap to obtain 95% confident intervals. Following(Excoffier *et al.* 2013) recommendation, the likelihood ratio G-statistics (*CLR* = log_10_(*CL_O_/CL_E_*), where *CL_O_* and *CL_E_* are the observed and estimated maximum composite likelihood, respectively) was computed to evaluate model goodness-of-fit. A non-significant *p*-value of this test indicates that the observed SFS is well explained by the model. In a second step, demographic inference was extended to all three spruce species using pairwise joint minor allele frequency spectra. For the three major domains of *P. abies*, parameter-values that were estimated during the first step were used (see Fig. 4 for a description of the model).

## Results

### Genetic diversity and population divergence

Genetic diversity was calculated at 0-fold and 4-fold sites (π_0_ and π_4_) and their ratio calculated for protein coding sequences of *P. omorika, P. obovata*, and the three main domains of *P. abies* (Alpine, Carpathian and Fennoscandian). On average, we used around 30,000 SNPs in 7,453 genes for each calculation. *P. omorika* harbored the lowest diversity (π_4_=0.0066), while *P. obovata* and *P. abies* exhibited larger values, ranging from 0.0072 to 0.0079. We found less difference in π0 values among the three species (0.0029~0.0032). This led to large π_0_/π_4_ ratios in all three species but especially in *P. omorika* (0.44 compared to 0.39 and 0.4, for *P. obovata* and *P. abies*, respectively, Table 1). Estimates of Tajimas’ *D* were slightly negative but not significantly different from zero for *P. obovata* (−0.176), for *P. abies* (−0.32~-0.42), and for their hybrid population in Kirov (−0.296). The positive value of Tajimas’ *D* observed in *P. omorika* (0.875) likely reflects the sudden and very recent population contraction experienced by the species.

Pairwise population fixation indices (*F_ST_*) in noncoding regions were highest between *P. omorika* and *P. abies* domains (average *Fst=* 0.66). The Fennoscandian domain was more closely related to *P. obovata* (*F_ST_* = 0.13) and to the hybrid population (*F_ST_* = 0.1) than the two other *P. abies* domains were *(Fst =* 0.15~0.22, Table 1). This lends further support to the population admixture or gene flow from *P. obovata* towards *P. abies* northern populations (Tsuda *et al.* 2016). Within *P. abies*, genetic distances ranged from 0.12 to 0.15, Alpine and Carpathian domains being the closest, and Alpine and Fennoscandian domains the farthest (Table 1).

**Table 1.** Genetic diversity (π x 10^-3^), Tajima’s *D*, and population divergence (*F_st_*).

| | $\pi_0$ | $\pi_4$ | $\pi_0/\pi_4$ | *D* | *D*<sub>coding</sub> | *P.omk* | *P.obv* | Kirov | FEN | ALP | CAR |
| --- | --- | --- | --- | --- | --- | --- | --- | --- | --- | --- | --- |
| *P. omk* | 2.9 | 6.6 | 0.44 | 0.88 | 0.04 | - | 0.642 | 0.639 | 0.652 | 0.677 | 0.671 |
| *P. obv* | 3.0 | 7.7 | 0.39 | -0.18 | -0.05 |  | - | 0.084 | 0.133 | 0.222 | 0.196 |
| *Kirov* | 3.9 | 10 | 0.39 | -0.30 | -0.16 |  |  | - | 0.102 | 0.172 | 0.147 |
| *FEN* | 3.2 | 7.9 | 0.41 | -0.34 | -0.22 |  |  |  | - | 0.145 | 0.123 |
| *ALP* | 2.7 | 7.2 | 0.38 | -0.32 | -0.22 |  |  |  |  | - | 0.116 |
| *CAR* | 2.9 | 7.3 | 0.40 | -0.42 | -0.26 |  |  |  |  |  | - |

*P. omk.*, *P. omorika*; *P. obv.*, *P. obovata*. *P. abies*: FEN, Fennoscandian, ALP, Alpine, and CAR, Carpathian.

### Population structure and admixture in P. abies

First, a principal component analysis (PCA) was conducted on genetic variation of *P. abies* and *P. obovata. P. omorika* was excluded from this analysis as its divergence from the two other species far exceeded the one between *P. abies* and *P. obovata* populations (Table 1 and 3). While *P. obovata* and the hybrid populations clustered separately from *P. abies*, the range of the latter also exhibited clear population genetic structure (Fig. 2a and b). Seven clusters can be distinguished within *P. abies* that are embedded within a triangle whose summits correspond to three main domains: Alpine, Carpathian and Fennoscandian. Other clusters appear as intermediate between these three domains: Central Europe between Carpathian and Alpine, Russian-Baltics between Carpathian and Fennoscandian and Southern and Central Sweden between the Alpine and Fennoscandian clusters. Trees from northern Poland constitute a separate cluster situated between the Russian-Baltics and the Carpathian domains.

**Figure 2.**
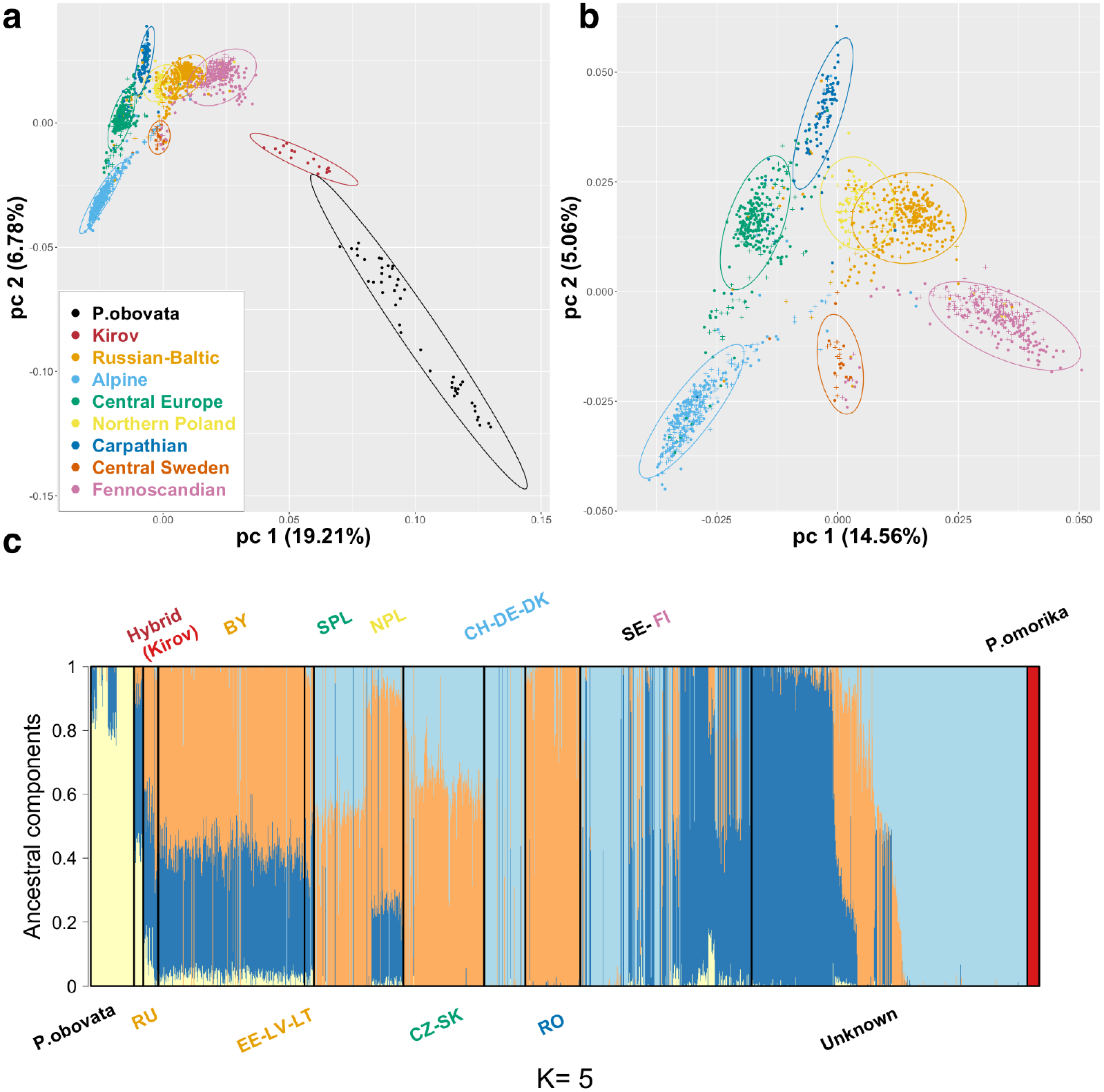
Population structure and admixture inferences of the three European spruces species. (a) Multidimensional scaling plot of PCA results (only *P. abies* and *obovata* individuals are represented), colors corresponds to different genetic clusters (see legend). The two first principal components are shown. (b) PCA results based on *P. abies* populations only. (c) Admixture analyses describing the proportion of ancestral components for K=5. Red and yellow are respectively, *P. omorika* and *P. obovata* genetic background, while light blue, orange and dark blue represent, Alpine, Carpathian, and Fennoscandian domains of *P. abies*, respectively. Solid black lines delimit the different populations. Trees with unclear origin in records are gathered under the “Unknown” label. Text colors showed the same genetic clusters in (a) except for the Swedish clusters. Russian-Baltic: Russia (RU), Belarus (BY), Estonia (EE), Latvia (LV), Lithuania (LT); Alpine: Germany (DE), Switzerland (CH), Denmark (DK); Central Europe: Slovakia (SK), Cze-republic (CZ), Southern Poland (SPL); Northern Poland (NPL); Carpathian: Romania (RO); Fennoscandia: Finland (FI), Sweden (SE). Note that all Swedish populations (SE) were mixed in this case.

To understand the history of the clusters detected in the PCA we turned to unsupervised clustering analysis (ADMIXTURE, Fig. 2c). Examination of the resulting Admixture plot led to the following interpretation. The three summits of the triangle in the PCA analysis, representing the Alpine, Carpathian and Fennoscandian domains correspond to three ancestral clusters in *P. abies* populations from which other clusters are derived through admixture. For instance, the Russian-Baltics populations include components from the Carpathians and Fennoscandia, as well as a small fraction from *P. obovata.* Central European populations derived from admixture between the Alpine and Carpathian domains. Trees from Northern Poland are similar to trees from the Russian-Baltics domain but also contain an additional contribution from the Alpine domain. The Swedish populations are particularly complex: some trees clustered with Alpine ones, others with Fennoscandia and the Russian-Baltics domain and a small fraction correspond to admixture between the Alpine and Fennoscandian domains.

Finally, no admixture was identified in *P. omorika* but traces of a Fennoscandian contribution could be detected in *P. obovata.* The populations Indigo and Kirov were sampled from the hybrid-zone at roughly the same longitude but at very different latitudes. Interestingly, only the southern population, Kirov, exhibits signs of admixture while the northern population Indigo belongs to *P. obovata* confirming that this species dominated further west at higher latitudes (Tsuda *et al.* 2016).

### Prediction of geographic origin based on genotype similarity

As noted above, information on the exact origin of 575 individuals of the breeding population was missing. We therefore used the PCA coordinates of these individuals to assign them to one of the seven *P. abies* clusters established from trees sampled in natural populations. Five-fold cross-validation showed that this method was robust (error rate < 8%). Each individual was assigned to one of the seven clusters identified in Norway spruce populations by PCA and a confidence value was computed by bootstrapping over 100 subsets of total samples. In total, 560 out of 575 (97.4%) of the individuals whose ancestry was unknown (“Unknown” individuals) could be assigned to one of the seven *P. abies* clusters with an average probability over 0.905 based on genetic similarity (Table 2), the 15 remaining belonging to *P. omorika.* Among the 560 trees that could be assigned, the two largest groups consisted of Alpine and Fennoscandian clusters (37% and 35%, respectively). The results thus suggest that the genomic markers we used in this study are powerful enough to give a high-resolution picture of the recent divergence history in Norway spruce. Hence, the method could serve as a rather efficient way to identify translocation material used for reforestation.

**Table 2.** Inference of geographic origin of *P. abies* trees based on genotype similarity.

| Cluster | Rus-Baltics | Alpine-Nordic |  |  | Visegrad* | North-Poland | Romania | Central-Sweden | Fennoscandia |
| --- | --- | --- | --- | --- | --- | --- | --- | --- | --- |
|  |  | Germany | Sweden | Denmark |  |  |  |  |  |
| #ind. | 38 | 27 | 168 | 13 | 53 | 28 | 17 | 21 | 195 |
| Prob. | 0.97 | 0.738 | 0.683 | 0.935 | 0.955 | 0.966 | 0.96 | 0.936 | 0.998 |

\*Visegrad includes Hungary, Poland, Czech Republic, and Slovakia

In addition, among the individuals that were sampled in common garden trials across central and southern Sweden, 290 showed a dominant Fennoscandian ancestry component, 379 displayed a dominant Alpine ancestry component that fit with postglacial re-colonization paths of *P. abies* (Tollefsrud *et al.* 2008; Tollefsrud *et al.* 2009) and thus could have truly originated from local populations and 31 shared both ancestries. The rest exhibited various levels of contribution from the Carpathian domain or even from *P. obovata.* This suggests that at least 55% of *P. abies* trees in Central and Southern Sweden are recent translocations and up to 75% when considering trees belonging to the Alpine domain as exogeneous.

Previous analyses revealed the importance of admixture event in *P. abies* recent history in Western Europe: TreeMix v1.13 (Pickrell and Pritchard 2012) was thus used to quantify the intensity and direction of the major admixture events between *Picea* populations (Fig. S2). They were (i) from *P. obovata* to the *P. abies-P. obovata* hybrids (41.5% of ancestry), (ii) from the hybrid to the Russia-Baltics cluster (39.0%), and (iii) from *P. obovata* to the Fennoscandian cluster (12.7%) (Fig. 3). All admixture events probably occurred quite recently as they were close to the tips of the tree (Fig. S2). Extensive admixture among *P. abies* populations was also identified using the *f_3_*-test, which is 3-population generalization of Fst that allows testing for admixture in a focal population (Reich *et al.* 2009). More than half of *f*_3_-statistics were significantly negative (56.5%) indicating admixed components in the Russia-Baltics individuals while 11.5% ~ 22% of f3-tests supported admixed components in Central Europe and in Fennoscandia populations (mainly from *P. obovata* and hybrid ancestry).

### Demography inference of ancestry populations in P. abies

Multidimensional SFS was first used to estimate the demographic history of the three most divergent *P. abies* clusters, Alpine (ALP), Carpathian (CAR), and Fennoscandian (FEN), which also represent the three main ancestry components of *P. abies* in the previous analyses. Assuming an average migration rate of 10^-6^, lead to similar estimates of Ne for the three *P. abies* domains, around 6,000 to 8,000 (95% CI: 5,000 – 11,400), with CAR and FEN populations slightly larger than ALP (Table 3 and Fig. 4). The ancestral effective population size of *P. abies* was much larger with an estimated value around 2.5×10^5^ to 5.7×10^5^. Assuming a generation time of 25 years and a mutation rate of 1.1×10^-9^ per site per year (Willyard *et al.* 2007; Chen *et al.* 2012b; Nystedt *et al.* 2013), the three populations had a similar divergence history since 15 million years ago (95% CI: 9 – 17.7 mya). More recently the populations went through a bottleneck around 13,000 years ago (95% CI: 6,400 – 33,000), which approximately corresponds to the end of LGM.

**Figure 4.**
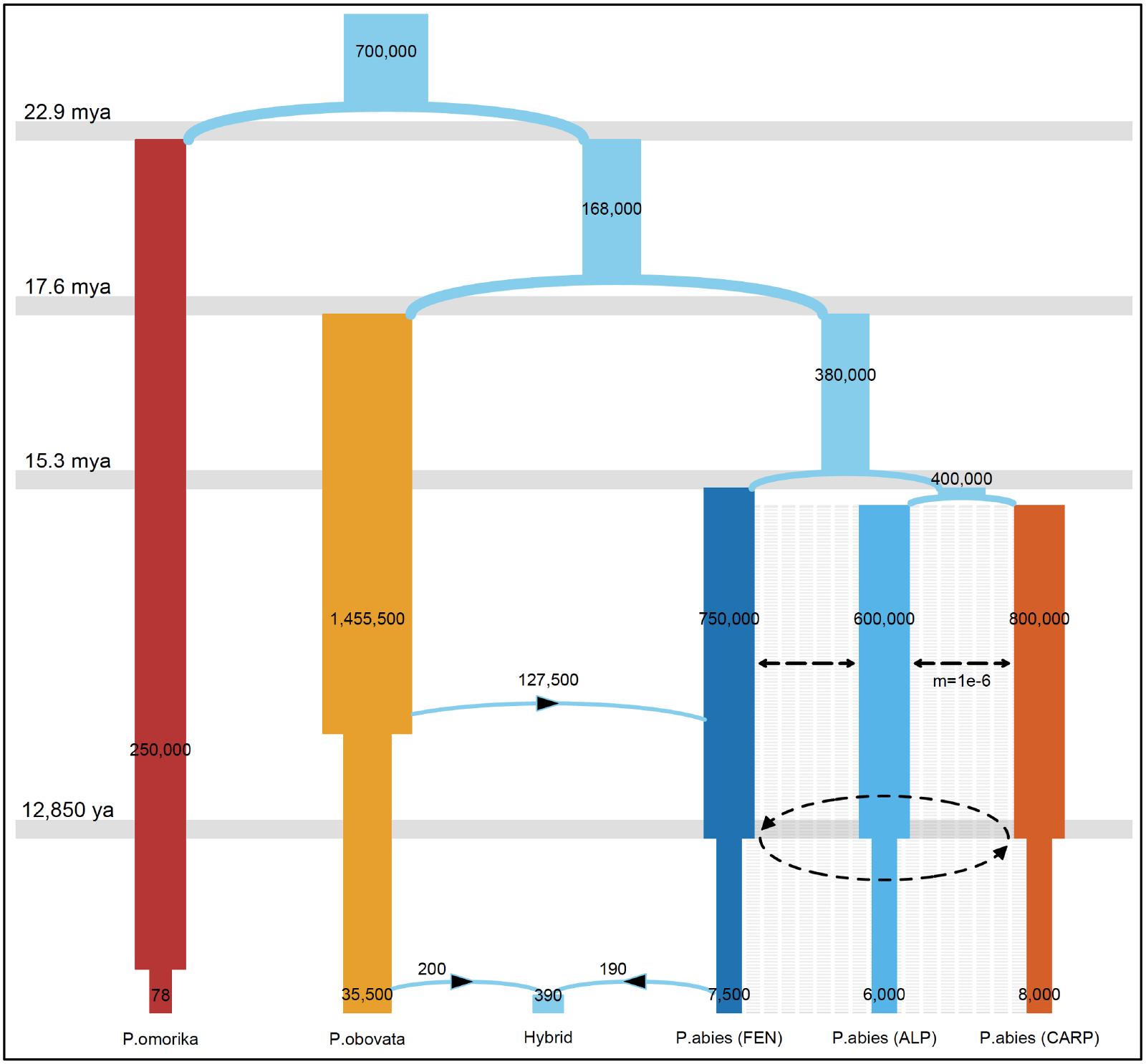
*Picea* genus demographic history in Europe. The best-fitting model describes bottlenecks for *P. omorika, P. obovata*, and *P. abies* (Alpine, Carpathian, and Fennoscandia domains) in different periods. The blue arrows describe the major admixture event from *P. obovata* to *P. abies* (FEN), and the hybridization between the two species. The black dotted line is migration and the figures within bars are the effective population sizes. Migration was allowed between the three *P. abies* domains with a fixed rate of 1×10^-6^. The parameters were estimated with FastSimCoal2.

**Table 3.**
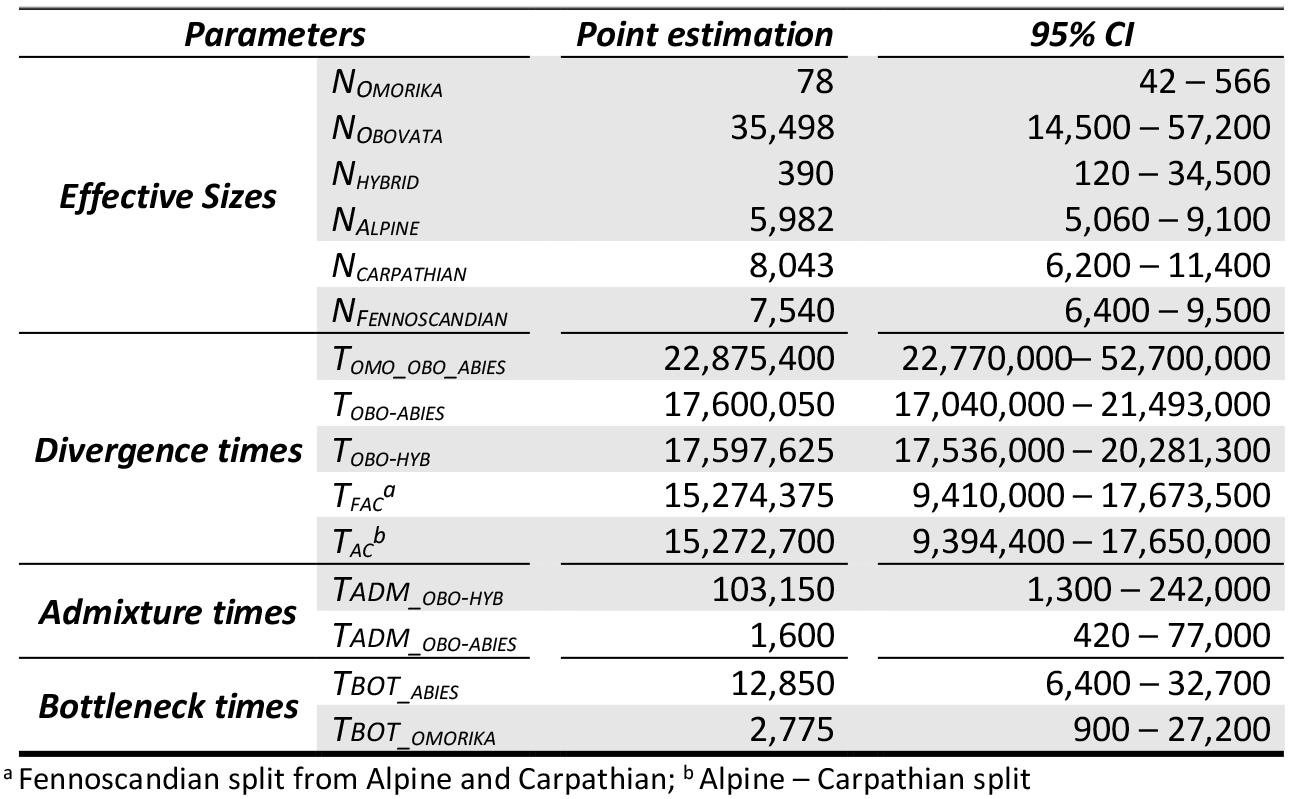
Demographic parameter estimatesfor *P. omorika (omo), P. obovata* (*obo*), *P. abies* main domains and *P. abies – P. obovata* hybrids (*hyb*).

Average migration rates (m) were set to 10^-6^ between the three main *P. abies* clusters. Given population sizes before bottleneck, it corresponded to approximately three migration events per generation per cluster. To assess if considering m > 0 improved the model likelihood significantly, composite likelihood parameter estimation was repeated with a null migration rate, m = 0. The Akaike’s weight of evidence supported a model with m > 0 in 95% of the runs. Demographic parameters were also inferred with free varying migration rate: the median migration rate was about 5.9×10^-6^. However, Akaike’s weight of evidence supported models considering a fixed migration parameter in 76 out of the 100 runs despite a slightly lower composite likelihood ratio (CLR) of models with migration free to vary.

The model goodness-of-fit was estimated by computing the likelihood ratio G-statistic test. For multidimensional model fitting, the p-value is highly significant which suggests that our model is certainly oversimplified and does not capture the full history of the species. However, likelihood ratio test based on a joint 2-dimensional SFS supports our divergence model (Fig. S3), indicating that, in spite of its simplified nature, it does capture the major features of the SFS (Anscombe residuals for pairwise joint SFS fitting are presented in Supplementary file S1).

### Divergence history of the three spruce species

We further extended our demographic inferences to all three spruce species and the hybrid between P. abies and P. obovata based on pairwise joint SFS. P. omorika has a particularly small effective population size (N0_omo<100), which is expected as the species is endangered and is part of the IUCN Red List. The natural range of P. omorika is believed to have been under continuing contraction since the LGM, but our estimates suggested a severe and recent bottleneck (~3200 fold) around 2,800 years ago. P. obovata has the largest Ne of the three species, with a value ~35,500. Unlike the other two species, no recent size contraction was detected in P. obovata after the end of LGM. The split of P. omorika occurred 23 mya, while the divergence between P. obovata and P. abies began 17.6 mya. Approximately 17% of P. abies FEN population migrated from P. obovata 103,000 years ago, when both species had much larger effective population sizes (6×10^5^ for FEN and 5.6×10^6^ for P. obovata). The hybrid population has a small effective population size around 400, and was established ~1,600 years ago, with almost equal contribution from both species (Fig. 4). The goodness-of-fit (GOF) for the model can be found in Supplementary file S2. We also provided re-scaled results to account for uncertainty about generation time (Table S1). With a 50-year generation time the estimates of effective populations size got halved but the divergence time in year remained the same.

## Discussion

In the present study we showed that the current distribution of genetic diversity in Norway spruce is the result of a complex mixture of ancient and recent demographic events and cannot be explained by relying on a simple phylogeographic paradigm. This history involves ancient splits, repeated hybridization events, severe bottlenecks, population movements, and very recently, important transfer of forest reproductive material associated with 19^th^ and 20^th^ centuries afforestation efforts. Below, we discuss the most salient features of our demographic inferences, starting with their impact on genetic diversity at synonymous and nonsynonymous sites.

### Sources of sequencing errors

There are two major possible sources of errors that could influence our inference on population admixture and demographics. First, our dataset was derived from coding regions or their flanking regions, whose allele frequencies could be affected by natural selection and thus would not be adequate for demographic inference. By calculating π_0_/π_4_ ratios at coding sequences, Chen *et al.* (2017) showed that the proportion of mutations that are putatively under weak purifying selection is non-negligible, especially for conifer trees. In the present study π_0_/π_4_ ratio is ~0.4 but Tajima’s D values tend to be small suggesting that purifying selection did not strongly affected the site frequency spectrum. Furthermore, linkage disequilibrium in spruce decays very fast within genes (within ~200 bp)(Heuertz *et al.* 2006; Chen *et al.* 2012a; Chen *et al.* 2014) so linked positive selection is also not likely to have affected nearby SNPs through hitchhiking or selective sweeps.

A second possible source of error comes from the fact that Norway spruce genome is incomplete and highly fragmented, which increases the risk of detecting false positive SNPs due to paralog sequences. To avoid this issue, we first aligned reads to the whole spruce genome instead of the baited sequences directly and chose for proper paired reads aligned to one unique position in the whole genome. Mapping to paralog sequences could also distort the distributions of quality scores. Instead of hard filtering based on arbitrary cutoffs, we applied a protocol of variant quality score recalibration, parameters of which were trained using set of SNPs discovered in a pedigree study (Baison *et al.* 2018; Bernhardsson *et al.* 2018). This should help reduce false positive discovery but may also bias towards shared SNPs and less private ones in *P. obovata* and *P. omorika.* It could result in overestimates of admixture or migration rates between *P. obovata* and *P. abies* and push back the divergence time among three species. However, the influence here is most likely minor since admixture is mainly in the hybrid and East European regions, which agrees with the conclusions of Tsuda *et al.* (2016) that was based on 10 SSR loci but with a much more extensive sampling across the ranges of *P. abies* and *P. obovata.* Furthermore, the distributions of variant quality scores for our SNP dataset also provided strong supports for our quality controls (Fig. S4).

### Genetic diversity

The π_0_/π_4_ is generally interpreted as a measure of the efficacy of purifying selection. However, it is also influenced by demography and, in particular, high values of π_0_/π_4_ can also result from the fact that the nonsynonymous diversity (π_0_) reaches equilibrium faster than synonymous diversity (π_4_), after a bottleneck (Gordo and Dionisio 2005, Brandvain and Wright 2016, Gravel (2016). In the present case the elevated π_0_/π_4_ ratio in *P. omorika* is mainly due to a decrease in π_4_ in that species (0.0066 compared to 0.0075 on average), due to the strong bottleneck experienced by the species as confirmed from our demographic simulations. Interestingly a similar decrease in genetic diversity was observed by (Kuittinen *et al.* 1991) when they compared variation at 19 allozyme loci in *P. omorika* and *P. abies* (expected heterozygosity *H_e_*=0.13-0.15 in *P. omorika* vs 0.22 in *P abies).* The estimates of synonymous diversity in the other two species were close to the value reported in interior spruce (0.0073) by Hodgins *et al.* (2016), however, with much smaller non-synonymous diversity (0.0013) in the latter. In the case of interior spruce the choice of conserved ortholog genes with lodgepole pine *(Pinus contorta)* may have led to strongly underestimate the genetic diversity at non-synonymous sites compared to synonymous sites. In general the π_0_/π_4_ ratio of spruce species is among the highest among perennial plants though it is still lower than in *Quercus robur* (~0.5) and other oak species (T. Leroy, pers. comm.)(Plomion *et al.* 2018). Possible explanations for the high π_0_/π_4_ ratio could be a more long-lasting effect of bottlenecks in long-lived organisms and/or a higher mutation rate in trees than in annual plants.

### Admixture and extensive translocation from central Europe to Southern Sweden

As in earlier studies, we identified three major genetic clusters (Alpine, Carpathian and Fennoscandian) in *P. abies* and extensive admixture from *P. obovata*, (Borghetti *et al.* 1988; Bucci and Vendramin 2000; Collignon *et al.* 2002; Achere *et al.* 2005; Heuertz *et al.* 2006; Tollefsrud *et al.* 2008; Tsuda *et al.* 2016; Fagernäs 2017). However, power provided by the large number of polymorphisms yielded a much more detailed and complex picture of the relationship among the Norway spruce main genetic clusters. In particular, admixture between the different clusters is much more frequent than suggested by previous studies, especially between the Alpine and Carpathian regions, and across the Baltic-Scandinavian regions. While the main thrust of the recolonization of previously glaciated areas after the last LGM, as well as previous waves of recolonization, stemmed primarily from the East there were also south-north movements as shown by a recent study of the vegetation in Poland during the Eemian Interglacial (130,000-115,000) and the periods that followed (Kupryjanowicz *et al.* 2018). During the Eemian Interglacial Kupryjanowicz *et al.* (2018) inferred a recolonization from the north-west and during the Holocene a recolonization of Poland from both the north-west and the south-east.

### Identifying recent transfers of material

The presence in central and southern Sweden, as well as Denmark, of trees with ancestry tracing back entirely, or almost entirely, to the Alpine or the Carpathian domain, reflects recent introductions. That these trees are recent introduction is supported by the fact that those individuals could be assigned to the same clade of Baltic and central European populations with very high confidence in both PCA and TreeMix analyses. Most likely, these genotypes with an important Alpine or Carpathian component reflect translocations from central European countries that took place during the massive reforestation of the twentieth century. Indeed, since the 1950s Sweden and Norway, and to a lesser extent Finland, started to import seeds for forest reproduction material from the Belarus, Czech Republic, Germany, Slovakia and the Baltic States (Myking *et al.* 2016). This is also documented in the Forestry Research Institute of Sweden (Skogforsk) records that indicate that 46 trees originated from German seedlots. Hence, our study illustrates that genomic markers could serve as an efficient way to identify and track the origins of translocations back to their origins. In addition, this study provides a prerequisite for investigating the genetic adaptive effects of translocation, which is of great importance for tree breeding, conservation, and forest management under climate change.

### Difference in demographic histories across spruce species

In contrast to other spruce species that had a negative Tajima’s D, *P. omorika’s* Tajima’s D was positive and relatively high, supporting the presence of a recent and sudden contraction of its population size. The most plausible demographic scenario for *P. omorika* indeed indicates a 3,000-fold reduction in population size around 2,800 years ago. This corresponds to the onset of the Iron Age and could have been followed by a severe depletion of forest resources although paleobotanical data from southwestern Bulgaria suggest that the onset of the Iron Age was associated with a reduction of *Abies alba* and *Pinus* sp. but a concomitant increase of *Fagus sylvatica* and *Picea abies* (Marinova *et al.* 2012). The species harbored before that time a rather stable and large population size, of the same order of magnitude to that of *P. abies.* The estimate of current effective population size is extremely small for *P. omorika*, which is consistent with the fact that a limited number of trees of *P. omorika* are distributed in small stands across Western Serbia and Eastern Bosnia and Herzegovina (Aleksić *et al.* 2017), whereas the other two spruce species are continentally distributed. More recently, a possible cause for the rapid decline of Serbian spruce could be a physiological stress induced by global warming that makes *P. omorika* more susceptible to *Armillaria ostoyae.* This pathogen is reported as a major cause for root rot and drying of the crown and the infected trees die within five to six years (Ivetić and Aleksić 2016; Aleksić *et al.* 2017). More generally, *P. omorika* could suffer from high temperatures. Gigov (1956) analysed pollen abundance in peat profiles at five locations across Serbia, and found that during the Subboreal period (5 – 2.5 ka BP), when the climate was continental, dry and warm, the percentage of *Picea cf. excelza* pollen declined at all studied localities, even at lower altitudes at which it was present during the previous Atlantic period (9-5 ka BP).

In contrast to the drastic change experienced by *P. omorika*, the three major domains of *P. abies* had similar effective population sizes, with the Alpine domain only slightly smaller. The three domains almost split at the same time and thereafter remained rather stable until the end of the GLM. *P. obovata* had a much larger population size and the bottleneck occurred earlier, around the end of the Eemian interglacial to the beginning of the last glacial. These results possibly reflect the fact that spruce species, as well as other tree species, especially in Siberia, actually occupied a much larger area before and during the LGM than assumed under the “southern refugia” paradigm that has dominated phylogeography until recently (Lascoux *et al.* 2004 and reference therein). The new picture that is emerging is one under which non-glaciated areas consisted in a tundra with scattered tree stands as found today at the tree line rather than an herbaceous tundra with trees only present in well delineated southern refugia. Trees might even have been able to survive in small pockets in glaciated areas (Tzedakis *et al.* 2013; Quinzin *et al.* 2017; Carcaillet *et al.* 2018).

Regarding divergence time, our result pushes the split of the three major domains in *P. abies* further back in time (15 mya), much older than the 5~6 mya from Lockwood *et al.* (2013); and Chen *et al.* (2016). A major reason could be that both latter estimates were based on admixed samples in either so-called “northern” or “southern” populations. The relatively recent admixture events between the three major domains could significantly reduce these estimates. While admixture could explain why previous estimates could be underestimates, the new divergence times nonetheless seem at variance with a median node age of less than 5 Mya obtained for northern clades by Leslie *et al.* (2012) although it should be pointed out that these estimates, in contrast, may as well err on the low side. For example in pines, a new fossil calibration led to an estimate of the origin of crown Pinus that is likely up to 30 Mya older (Early Cretaceous) than inferred in most previous studies (Late Cretaceous) (Saladin *et al.* 2017). We also estimated the split of *P. abies* and *P. obovata* to be around 18 mya and the split with *P. omorika* to be around 23 mya. This latter estimate is consistent with a divergence of the Eurasian spruce clade around the early to the middle Miocene (13~23 mya), as inferred from molecular clock dating methods (Leslie *et al.* 2012; Lockwood *et al.* 2013).

## Conclusion

Two main conclusions emerge from the present work. First, the current distribution of *P. abies* is the result of complex population movements and admixture events with at least three major ancestral groups (Alpine, Carpathian and Fennoscandia) and derived ones. As in Tsuda *et al.* (2016) we observed that admixture of *P. obovata* extended as far west as Fennoscandia and the Baltics.

Second, a large proportion of the trees selected in Southern Sweden to establish the Swedish breeding program were shown here to be recent introductions from Central Europe. This agrees well with previous studies of the release of “alien” material in Sweden (Laikre *et al.* 2006; Myking *et al.* 2016) that reported a large use of material of foreign origin in Swedish forestry. The trees genotyped in the present study were “plus trees” used to establish the breeding program and were therefore selected based on their superior phenotypes. Hence, it already suggests that the use of alien material did not have any strong negative effect on growth and productivity. In addition, since these trees have been tested through a large number of progeny or clonal tests across Southern Sweden this material provides a unique opportunity to test for local adaptation in Norway spruce and this is the object of a companion paper (Milesi *et al.* 2018).

## Data accessibility

Raw reads are available in SRA NCBI database (https://www.ncbi.nlm.nih.gov/sra/) deposited as PRJNA511374 bio-project. Scripts and datasets are deposited in Zenodo database (https://zenodo.org/) under DOI: 10.5281/zenodo.2530736.

## Supporting Information

**Table S1:** Demographic parameter estimates rescaled by a generation time of 50 years.

**Fig. S1:** Cross-validation error for unsupervised population clustering.

**Fig. S2:** TreeMix Graph of migration events in *P. abies* genus.

**Fig. S3:** Likelihood ratio G-statistics distribution.

**Fig. S4:** Variant quality scores reported for final SNP dataset.

**Supplementary files 1 and 2:** Anscombe residuals for pairwise joint SFS.

## Acknowledgements

The present study was financed by the Swedish Research Council for Environment, Agricultural Sciences and Spatial Planning (FORMAS) to ML, GJ and BK. JC was financially supported by the Swedish Foundation for Strategic Research (SSF) project “Genomic selection of Norway spruce for new bioproducts”. GGV was financially supported by the European Commission through the Dynabeech project (5th Framework Programme, QLRT-1999-01210). This preprint has been reviewed and recommended by Peer Community In Evolutionary Biology (https://dx.doi.org/10.24072/pci.evolbiol.100064).

## Conflict of interest disclosure

The authors of this preprint declare that they have no financial conflict of interest with the content of this article. Martin Lascoux is recommender at *PCI Evol. Biol.*

