## Supplementary figures and images for "Genomic data provides new insights on the demographic history and the extent of recent material transfers in Norway spruce"

### Supplementary file 1

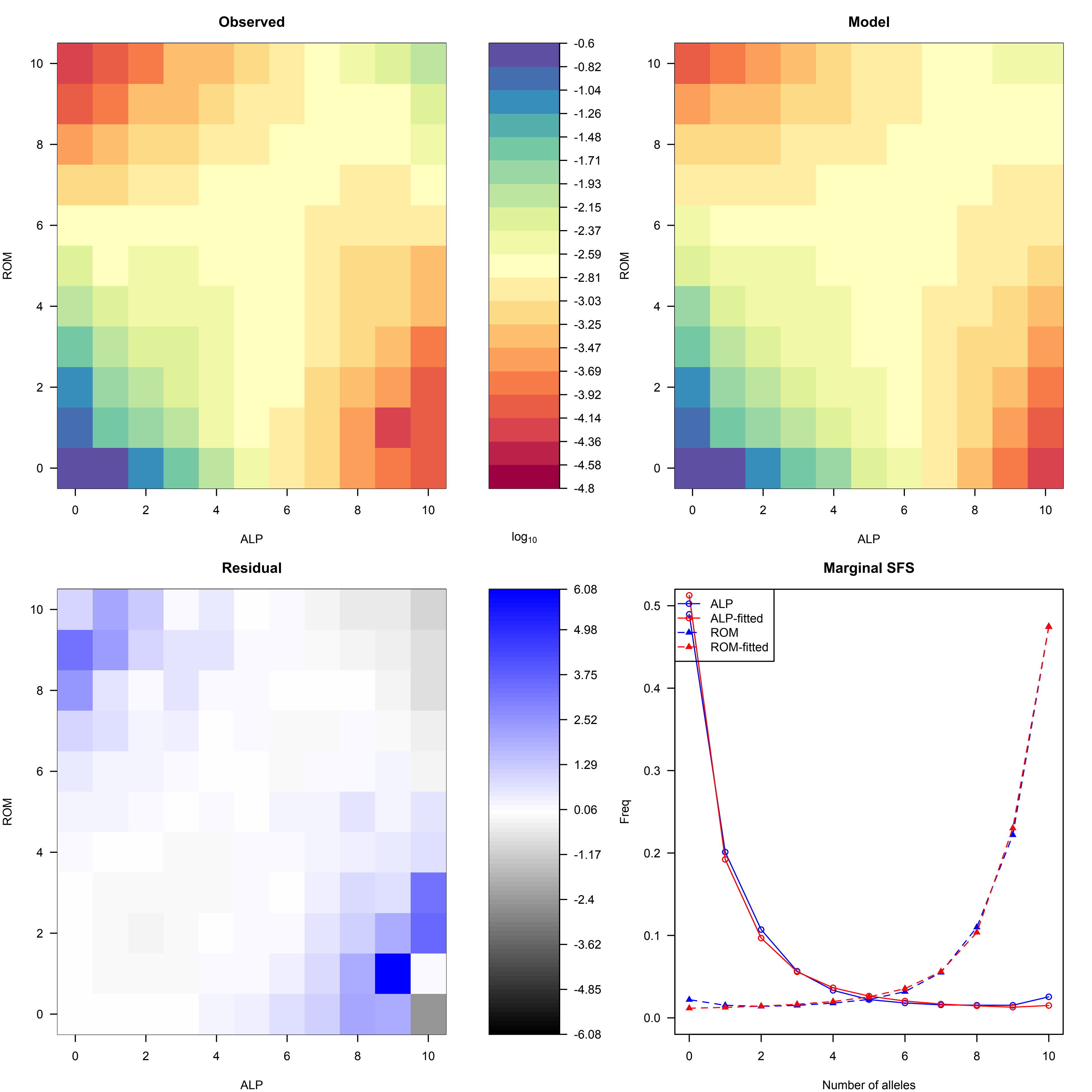

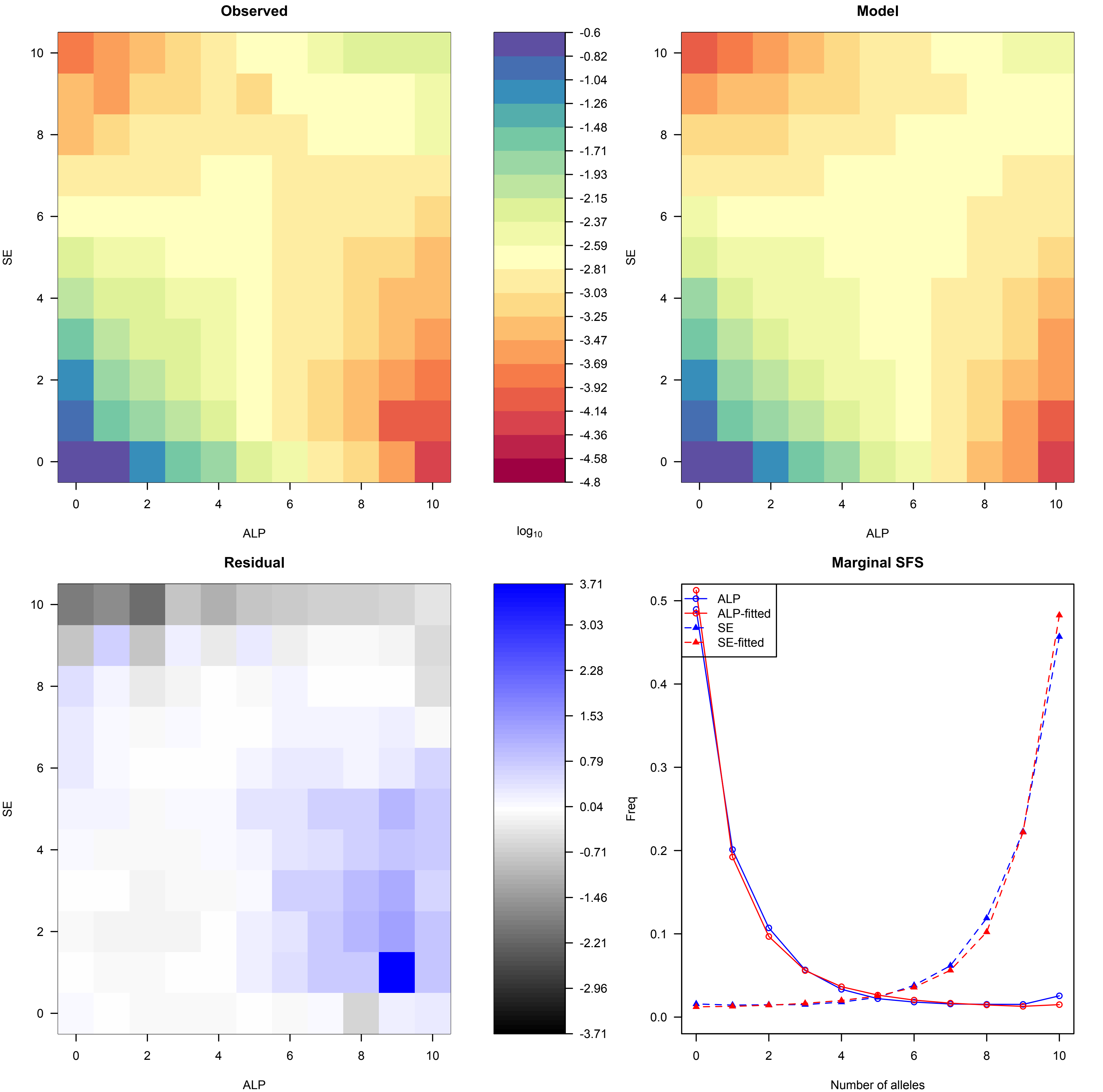

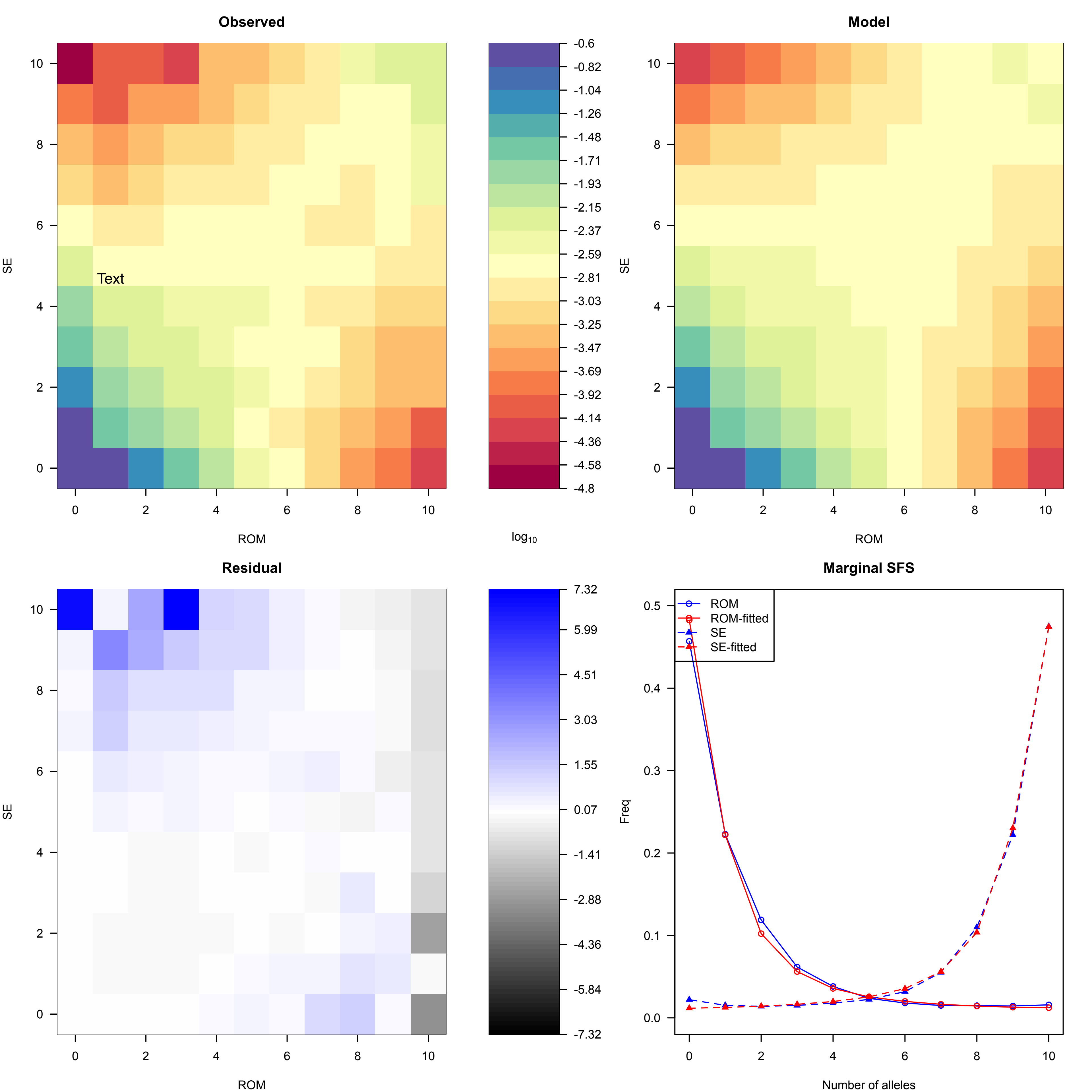
