## Supplementary file 2 for "Genomic data provides new insights on the demographic history and the extent of recent material transfers in Norway spruce"

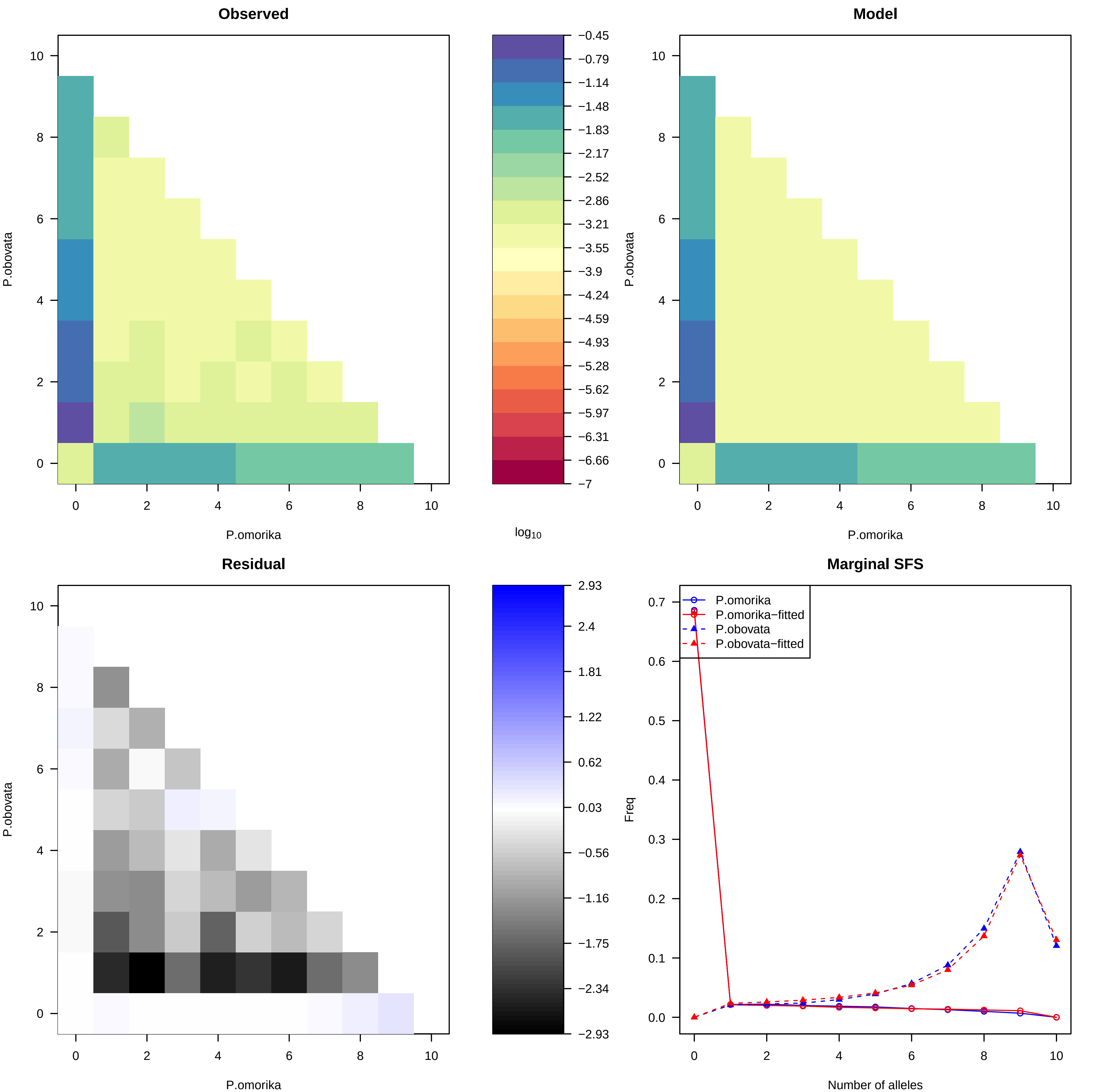

Observed
Model
P.obovata
P.omorika
log10
−0.45
−0.79
−1.14
−1.48
−1.83
−2.17
−2.52
−2.86
−3.21
−3.55
−3.9
−4.24
−4.59
−4.93
−5.28
−5.62
−5.97
−6.31
−6.66
−7
Residual
2.93
2.4
1.81
1.22
0.62
0.03
−0.56
−1.16
−1.75
−2.34
−2.93
Marginal SFS
P.omorika
P.omorika−fitted
P.obovata
P.obovata−fitted
Freq
Number of alleles

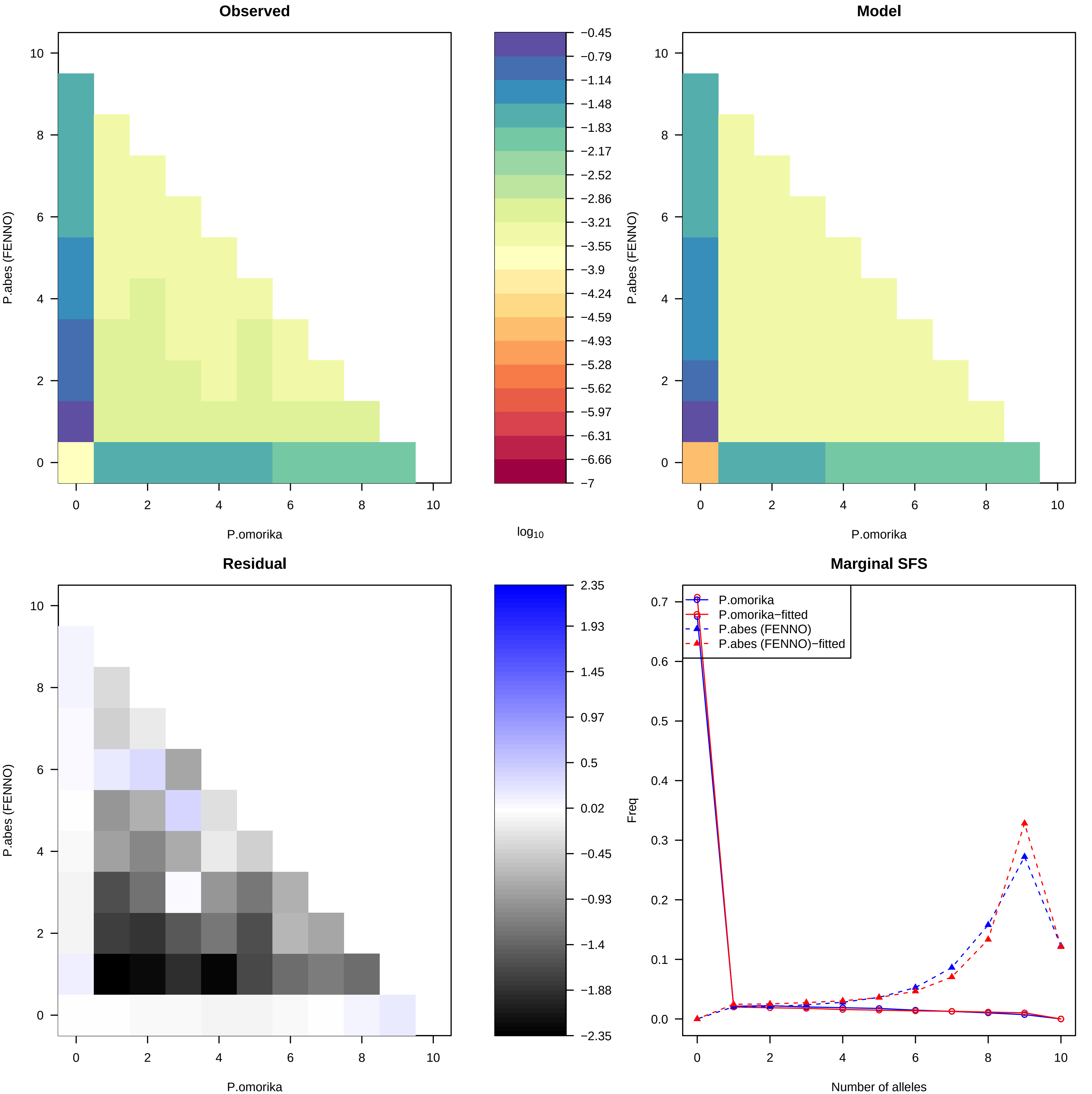

Observed
Model
Residual
Marginal SFS
P.abes (FENNO)
P.omorika
log10
Freq
Number of alleles
P.omorika
P.omorika−fitted
P.abes (FENNO)
P.abes (FENNO)−fitted
−0.45
−0.79
−1.14
−1.48
−1.83
−2.17
−2.52
−2.86
−3.21
−3.55
−3.9
−4.24
−4.59
−4.93
−5.28
−5.62
−5.97
−6.31
−6.66
−7
2.35
1.93
1.45
0.97
0.5
0.02
−0.45
−0.93
−1.4
−1.88
−2.35

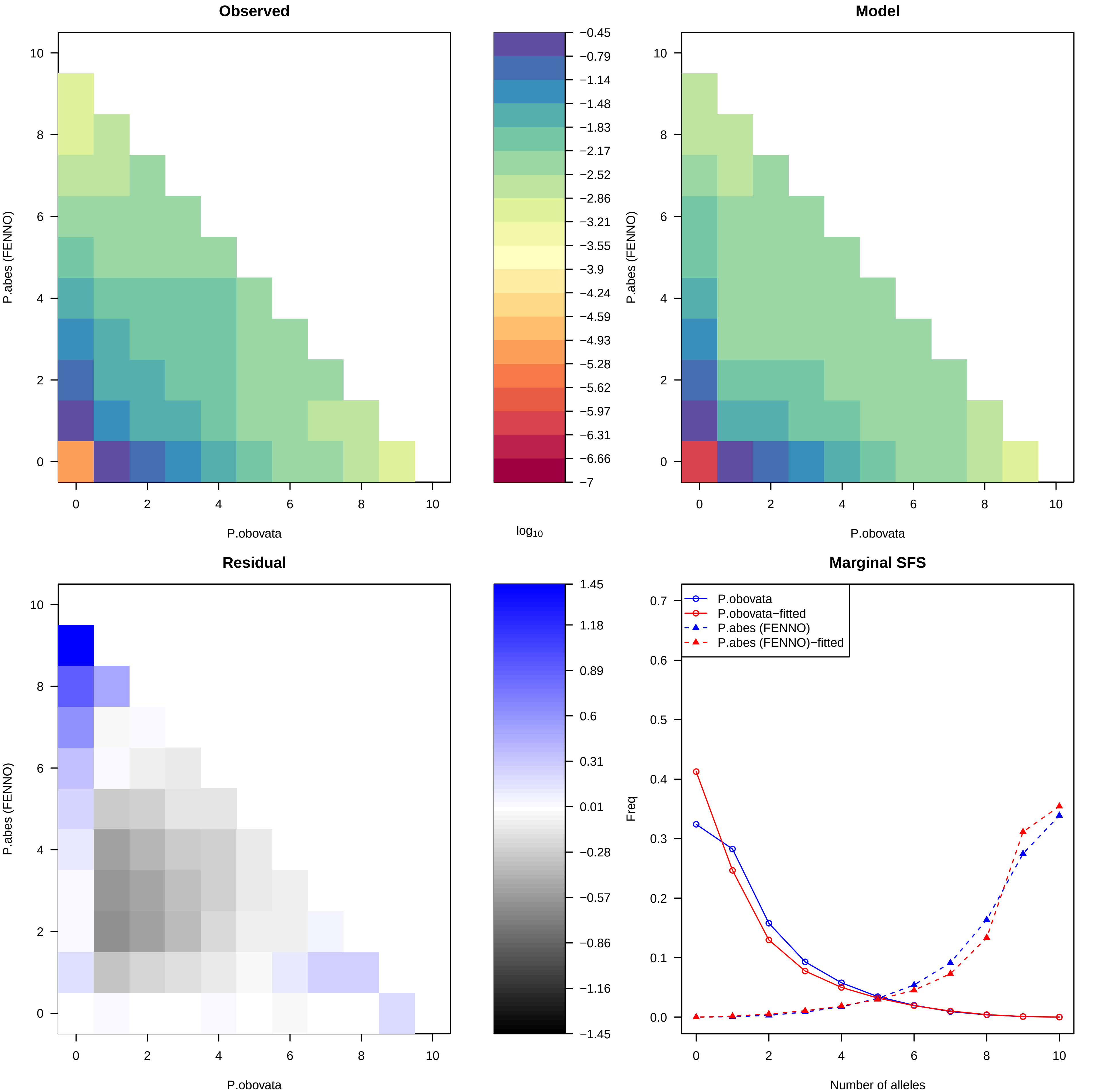

Observed
Model
Residual
Marginal SFS
P.abes (FENNO)
P.obovata
log10
−0.45
−0.79
−1.14
−1.48
−1.83
−2.17
−2.52
−2.86
−3.21
−3.55
−3.9
−4.24
−4.59
−4.93
−5.28
−5.62
−5.97
−6.31
−6.66
−7
1.45
1.18
0.89
0.6
0.31
0.01
−0.28
−0.57
−0.86
−1.16
−1.45
Freq
Number of alleles
P.obovata
P.obovata−fitted
P.abes (FENNO)
P.abes (FENNO)−fitted

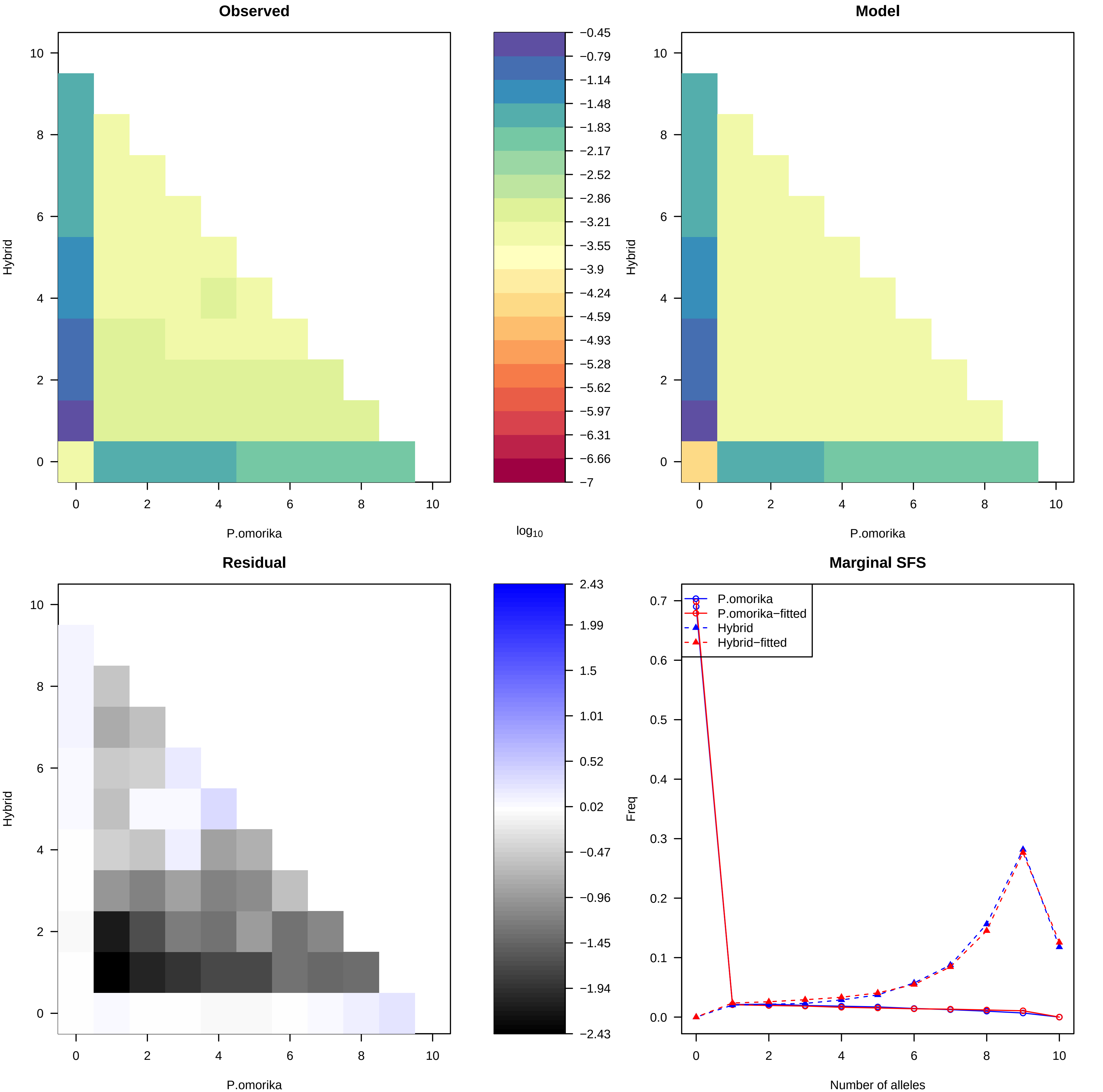
Observed
Model
Residual
Marginal SFS
Hybrid
P.omorika
log10
−0.45
−0.79
−1.14
−1.48
−1.83
−2.17
−2.52
−2.86
−3.21
−3.55
−3.9
−4.24
−4.59
−4.93
−5.28
−5.62
−5.97
−6.31
−6.66
−7
2.43
1.99
1.5
1.01
0.52
0.02
−0.47
−0.96
−1.45
−1.94
−2.43
P.omorika
P.omorika−fitted
Hybrid
Hybrid−fitted
Freq
Number of alleles

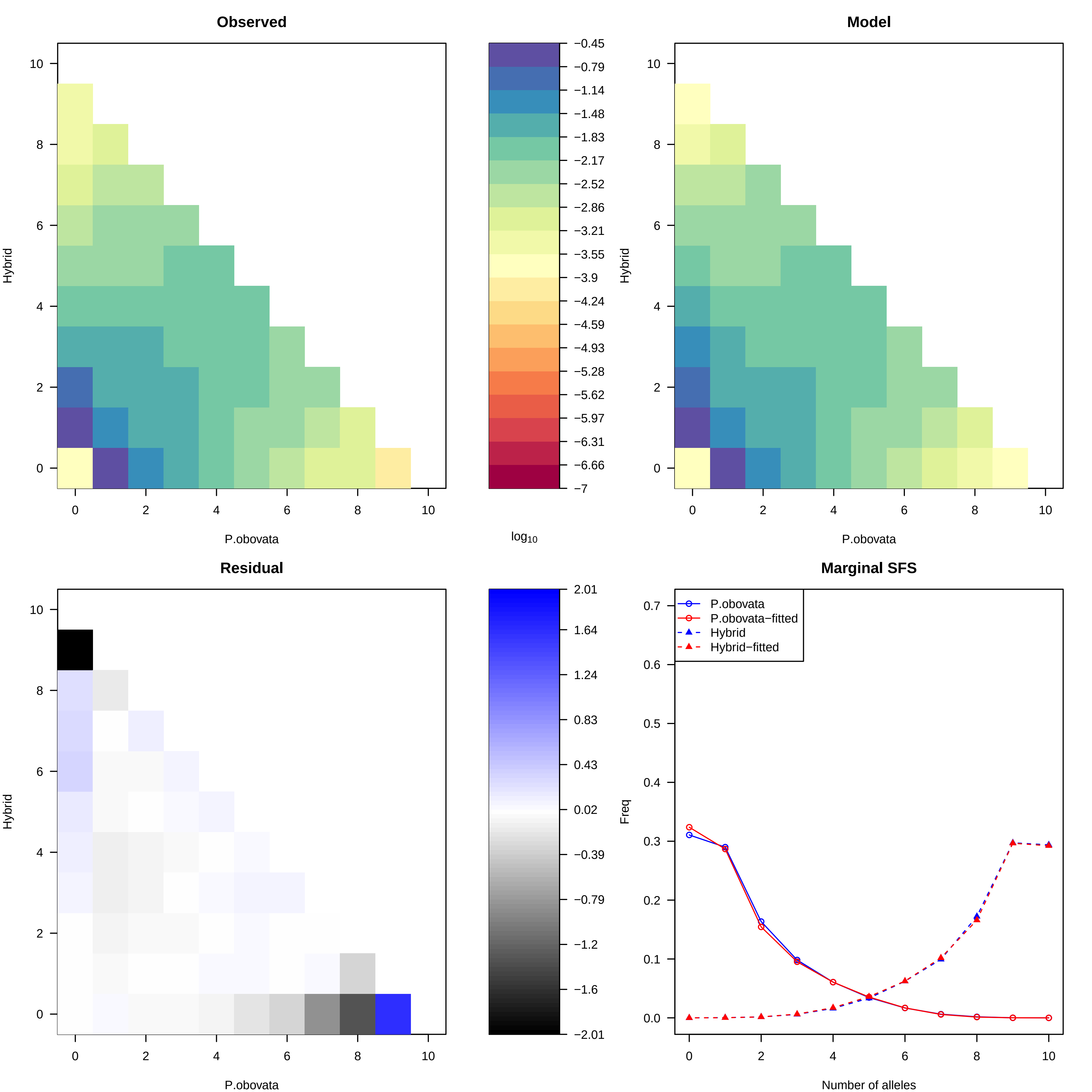

Observed
Model
Residual
Marginal SFS
Hybrid
P.obovata
log10
Freq
Number of alleles
P.obovata
P.obovata−fitted
Hybrid
Hybrid−fitted
−0.45
−0.79
−1.14
−1.48
−1.83
−2.17
−2.52
−2.86
−3.21
−3.55
−3.9
−4.24
−4.59
−4.93
−5.28
−5.62
−5.97
−6.31
−6.66
−7
2.01
1.64
1.24
0.83
0.43
0.02
−0.39
−0.79
−1.2
−1.6
−2.01

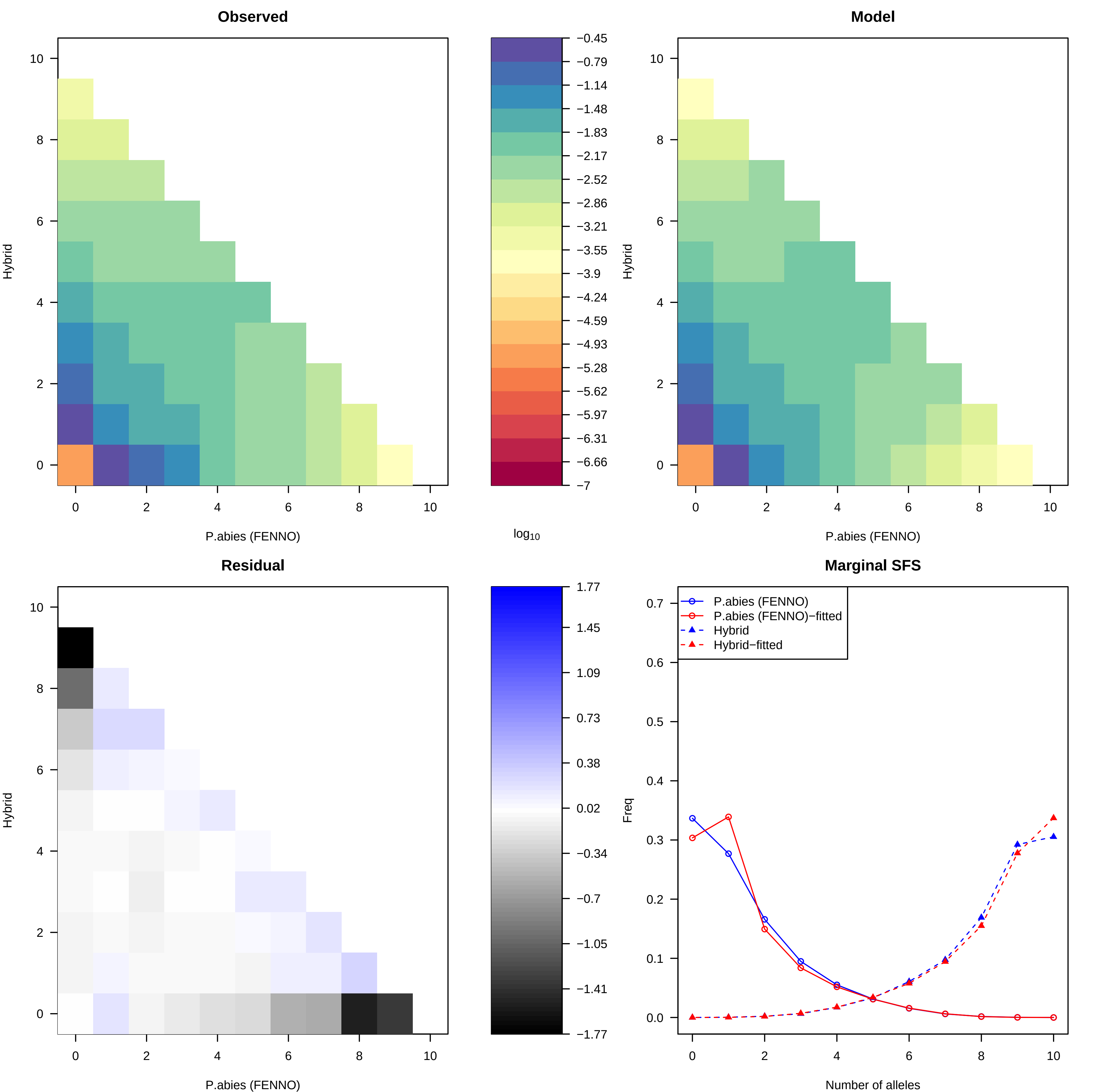

Observed
Model
Residual
Marginal SFS
Hybrid
P.abies (FENNO)
log10
−0.45
−0.79
−1.14
−1.48
−1.83
−2.17
−2.52
−2.86
−3.21
−3.55
−3.9
−4.24
−4.59
−4.93
−5.28
−5.62
−5.97
−6.31
−6.66
−7
1.77
1.45
1.09
0.73
0.38
0.02
−0.34
−0.7
−1.05
−1.41
−1.77
P.abies (FENNO)
P.abies (FENNO)−fitted
Hybrid
Hybrid−fitted
Freq
Number of alleles

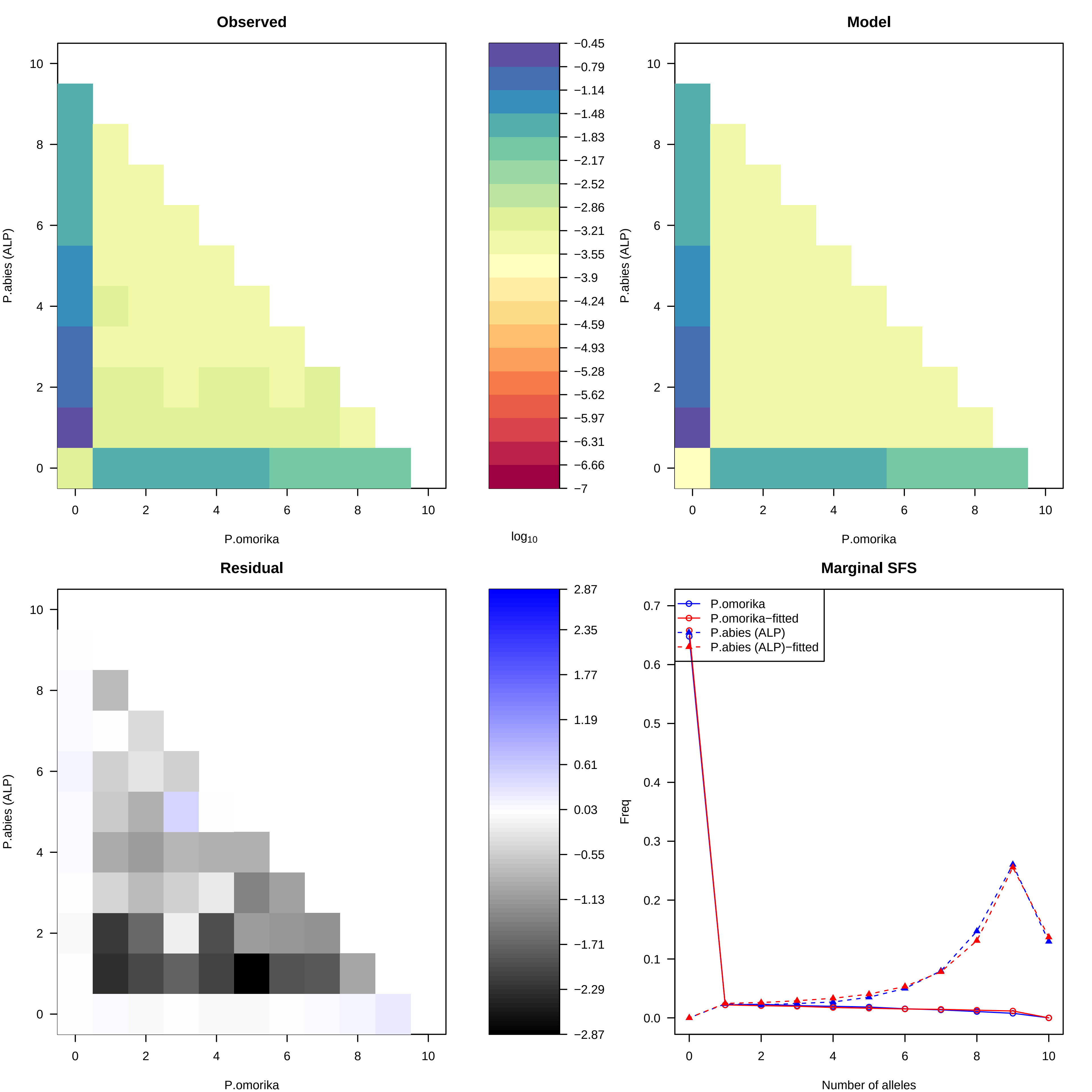
Observed
Model
Residual
Marginal SFS
P.abies (ALP)
P.omorika
log10
Freq
Number of alleles
P.omorika
P.omorika−fitted
P.abies (ALP)
P.abies (ALP)−fitted
−0.45
−0.79
−1.14
−1.48
−1.83
−2.17
−2.52
−2.86
−3.21
−3.55
−3.9
−4.24
−4.59
−4.93
−5.28
−5.62
−5.97
−6.31
−6.66
−7
2.87
2.35
1.77
1.19
0.61
0.03
−0.55
−1.13
−1.71
−2.29
−2.87

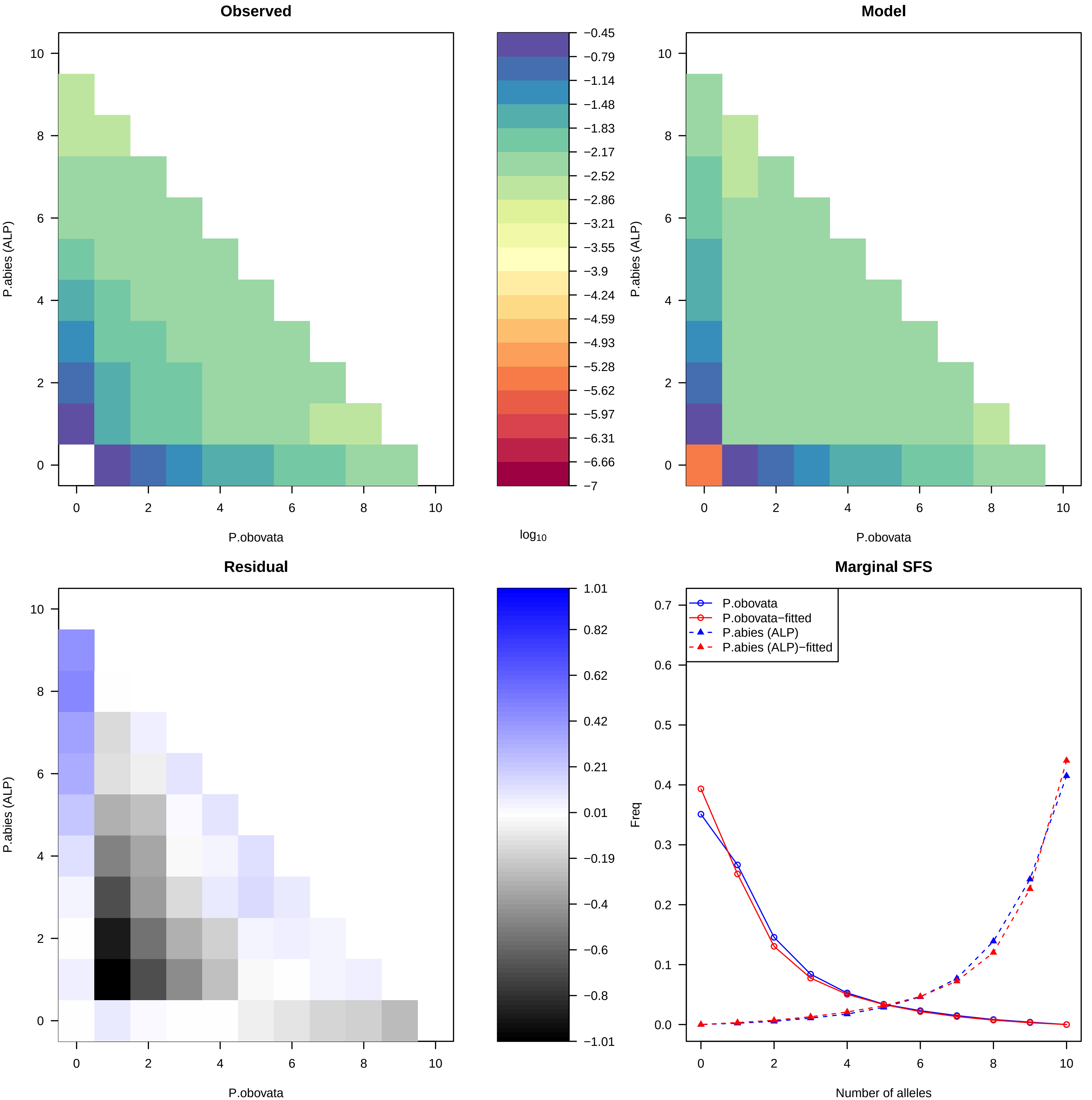

Observed
Model
Residual
Marginal SFS
P.abies (ALP)
P.obovata
log10
Freq
Number of alleles
P.obovata
P.obovata−fitted
P.abies (ALP)
P.abies (ALP)−fitted
−0.45
−0.79
−1.14
−1.48
−1.83
−2.17
−2.52
−2.86
−3.21
−3.55
−3.9
−4.24
−4.59
−4.93
−5.28
−5.62
−5.97
−6.31
−6.66
−7
1.01
0.82
0.62
0.42
0.21
0.01
−0.19
−0.4
−0.6
−0.8
−1.01

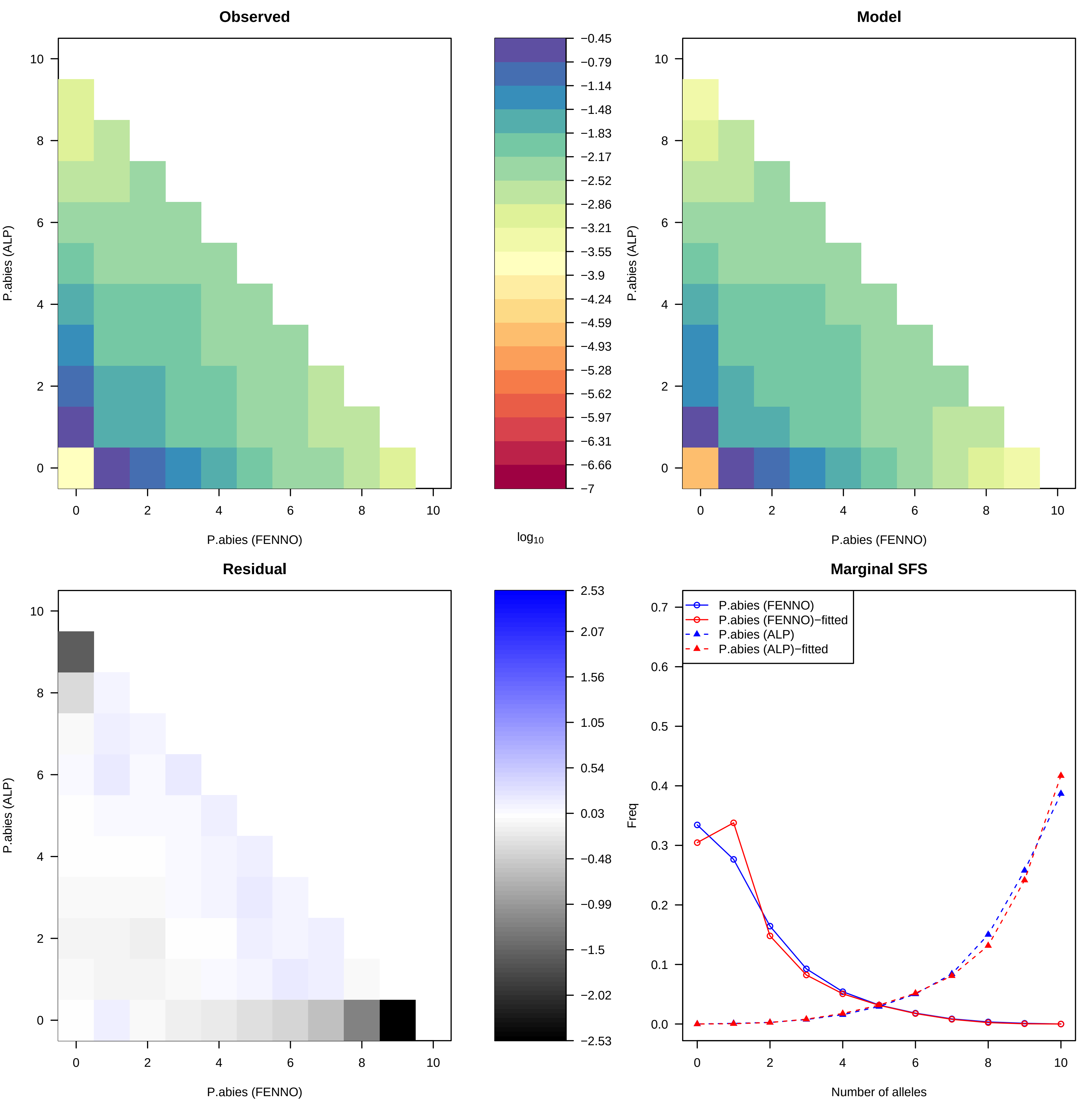

Observed
Model
Residual
Marginal SFS
P.abies (FENNO)
P.abies (ALP)
log10
−0.45
−0.79
−1.14
−1.48
−1.83
−2.17
−2.52
−2.86
−3.21
−3.55
−3.9
−4.24
−4.59
−4.93
−5.28
−5.62
−5.97
−6.31
−6.66
−7
2.53
2.07
1.56
1.05
0.54
0.03
−0.48
−0.99
−1.5
−2.02
−2.53
P.abies (FENNO)
P.abies (FENNO)−fitted
P.abies (ALP)
P.abies (ALP)−fitted
Freq
Number of alleles

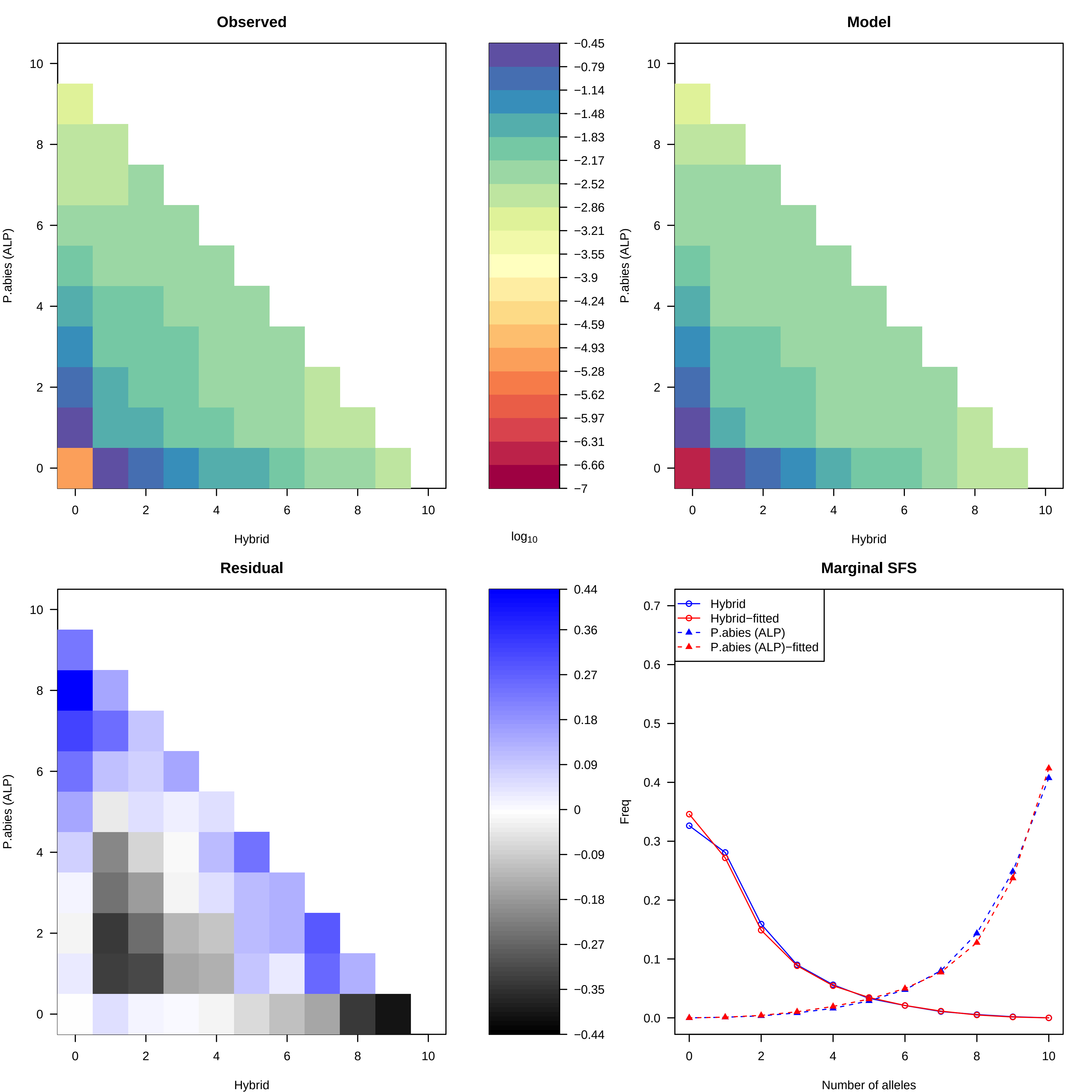

Observed
Model
Residual
Marginal SFS
P.abies (ALP)
Hybrid
log10
Freq
Number of alleles
Hybrid
Hybrid−fitted
P.abies (ALP)
P.abies (ALP)−fitted
−0.45
−0.79
−1.14
−1.48
−1.83
−2.17
−2.52
−2.86
−3.21
−3.55
−3.9
−4.24
−4.59
−4.93
−5.28
−5.62
−5.97
−6.31
−6.66
−7
0.44
0.36
0.27
0.18
0.09
0
−0.09
−0.18
−0.27
−0.35
−0.44

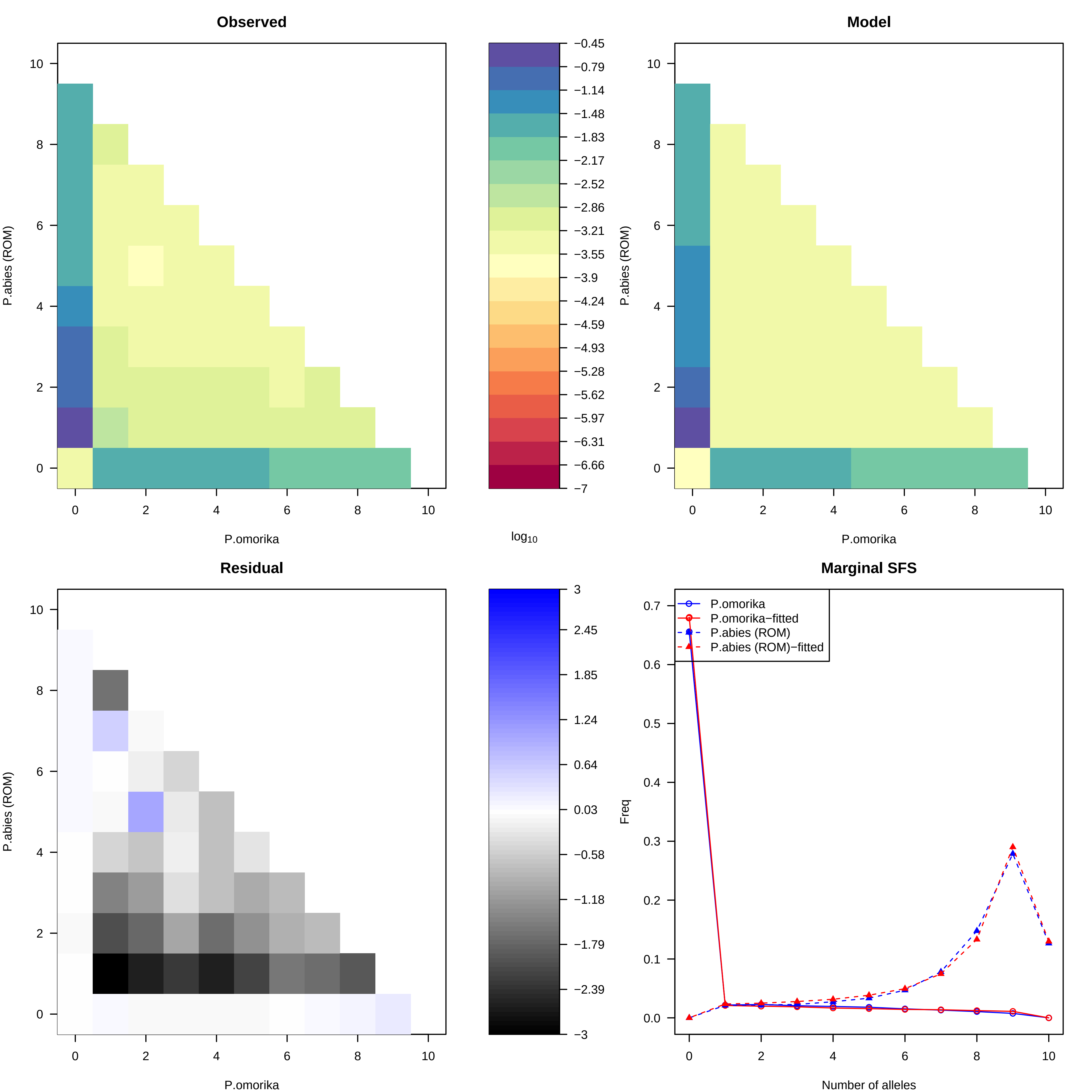
Observed
Model
Residual
Marginal SFS
P.abies (ROM)
P.omorika
log10
Freq
Number of alleles
P.omorika
P.omorika−fitted
P.abies (ROM)
P.abies (ROM)−fitted
−0.45
−0.79
−1.14
−1.48
−1.83
−2.17
−2.52
−2.86
−3.21
−3.55
−3.9
−4.24
−4.59
−4.93
−5.28
−5.62
−5.97
−6.31
−6.66
−7
3
2.45
1.85
1.24
0.64
0.03
−0.58
−1.18
−1.79
−2.39
−3

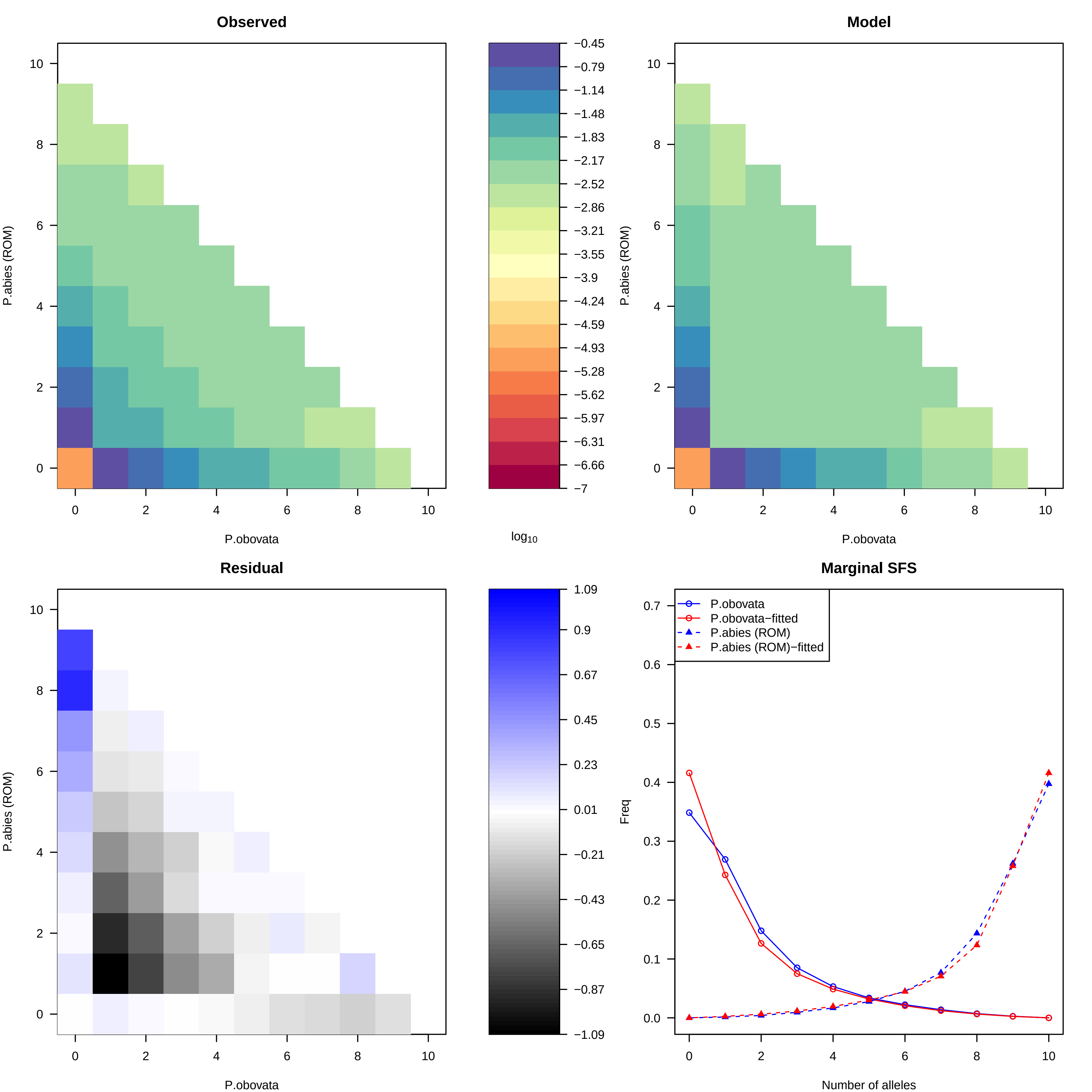

Observed
Model
Residual
Marginal SFS
P.abies (ROM)
P.obovata
log10
−0.45
−0.79
−1.14
−1.48
−1.83
−2.17
−2.52
−2.86
−3.21
−3.55
−3.9
−4.24
−4.59
−4.93
−5.28
−5.62
−5.97
−6.31
−6.66
−7
1.09
0.9
0.67
0.45
0.23
0.01
−0.21
−0.43
−0.65
−0.87
−1.09
Freq
Number of alleles
P.obovata
P.obovata−fitted
P.abies (ROM)
P.abies (ROM)−fitted

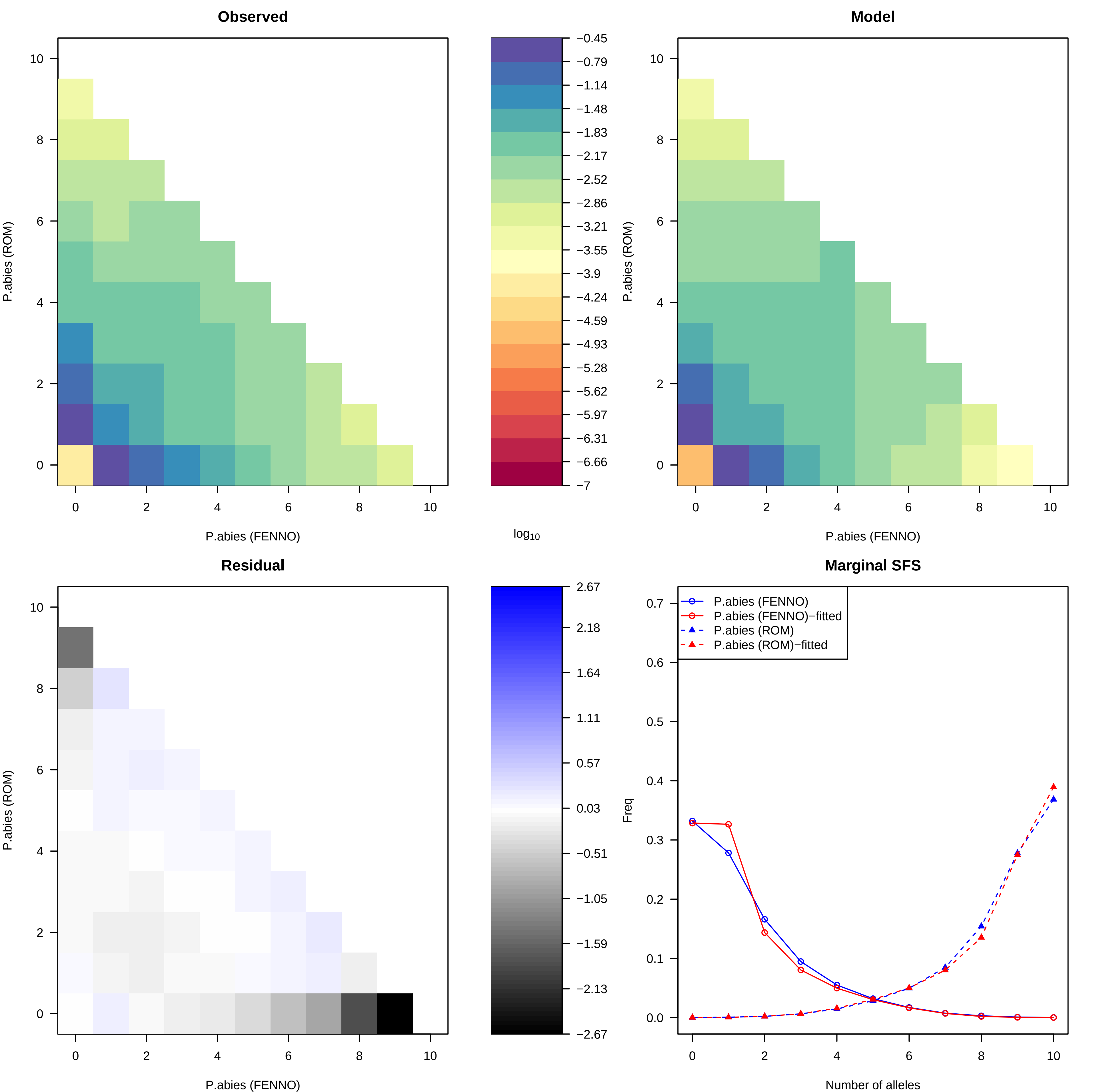
Observed
Model
Residual
Marginal SFS
P.abies (ROM)
P.abies (FENNO)
log10
−0.45
−0.79
−1.14
−1.48
−1.83
−2.17
−2.52
−2.86
−3.21
−3.55
−3.9
−4.24
−4.59
−4.93
−5.28
−5.62
−5.97
−6.31
−6.66
−7
2.67
2.18
1.64
1.11
0.57
0.03
−0.51
−1.05
−1.59
−2.13
−2.67
P.abies (FENNO)
P.abies (FENNO)−fitted
P.abies (ROM)
P.abies (ROM)−fitted
Freq
Number of alleles

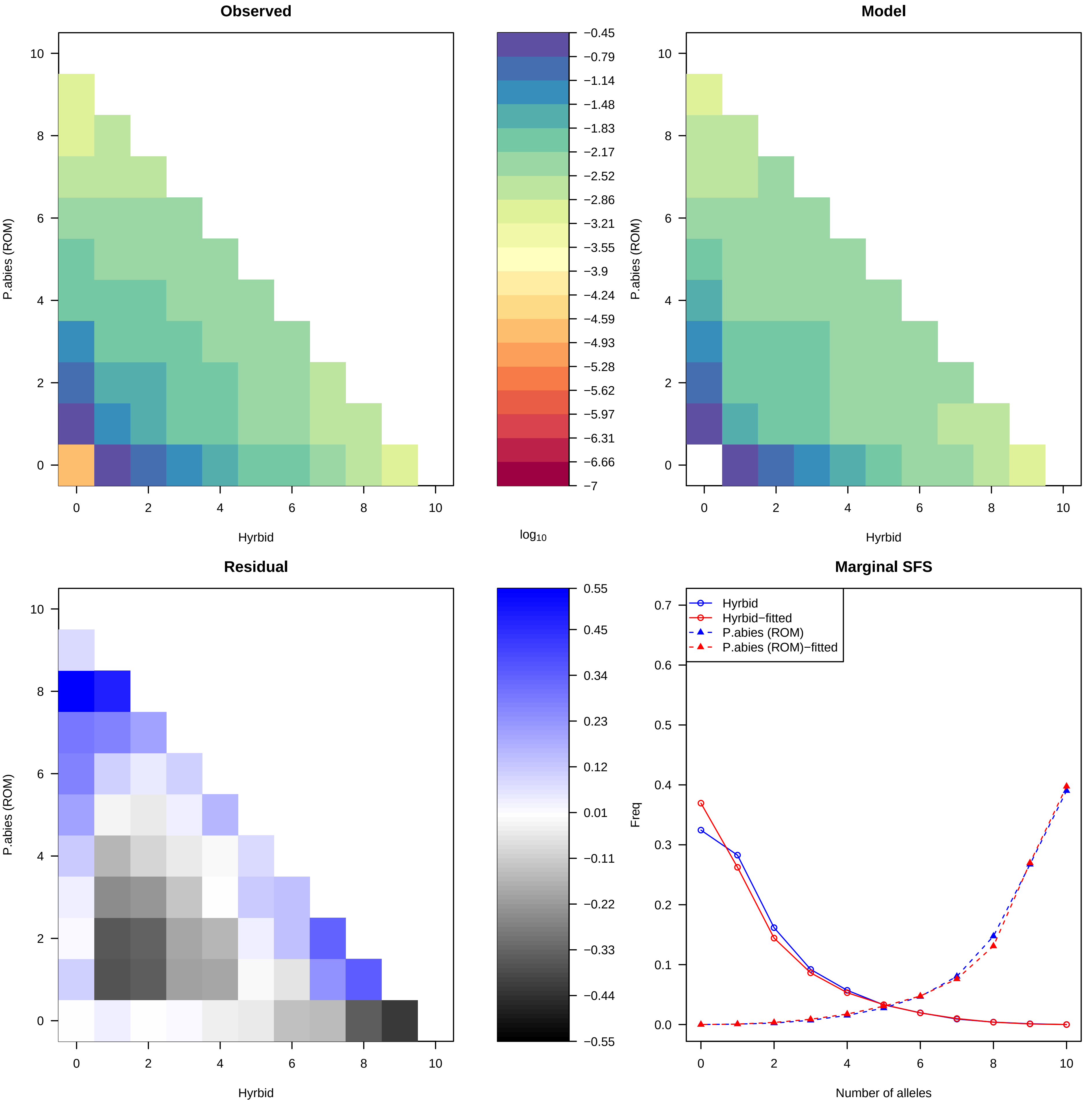
Observed
Model
Residual
Marginal SFS
P.abies (ROM)
Hyrbid
log10
−0.45
−0.79
−1.14
−1.48
−1.83
−2.17
−2.52
−2.86
−3.21
−3.55
−3.9
−4.24
−4.59
−4.93
−5.28
−5.62
−5.97
−6.31
−6.66
−7
0.55
0.45
0.34
0.23
0.12
0.01
−0.11
−0.22
−0.33
−0.44
−0.55
Hyrbid
Hyrbid−fitted
P.abies (ROM)
P.abies (ROM)−fitted
Freq
Number of alleles

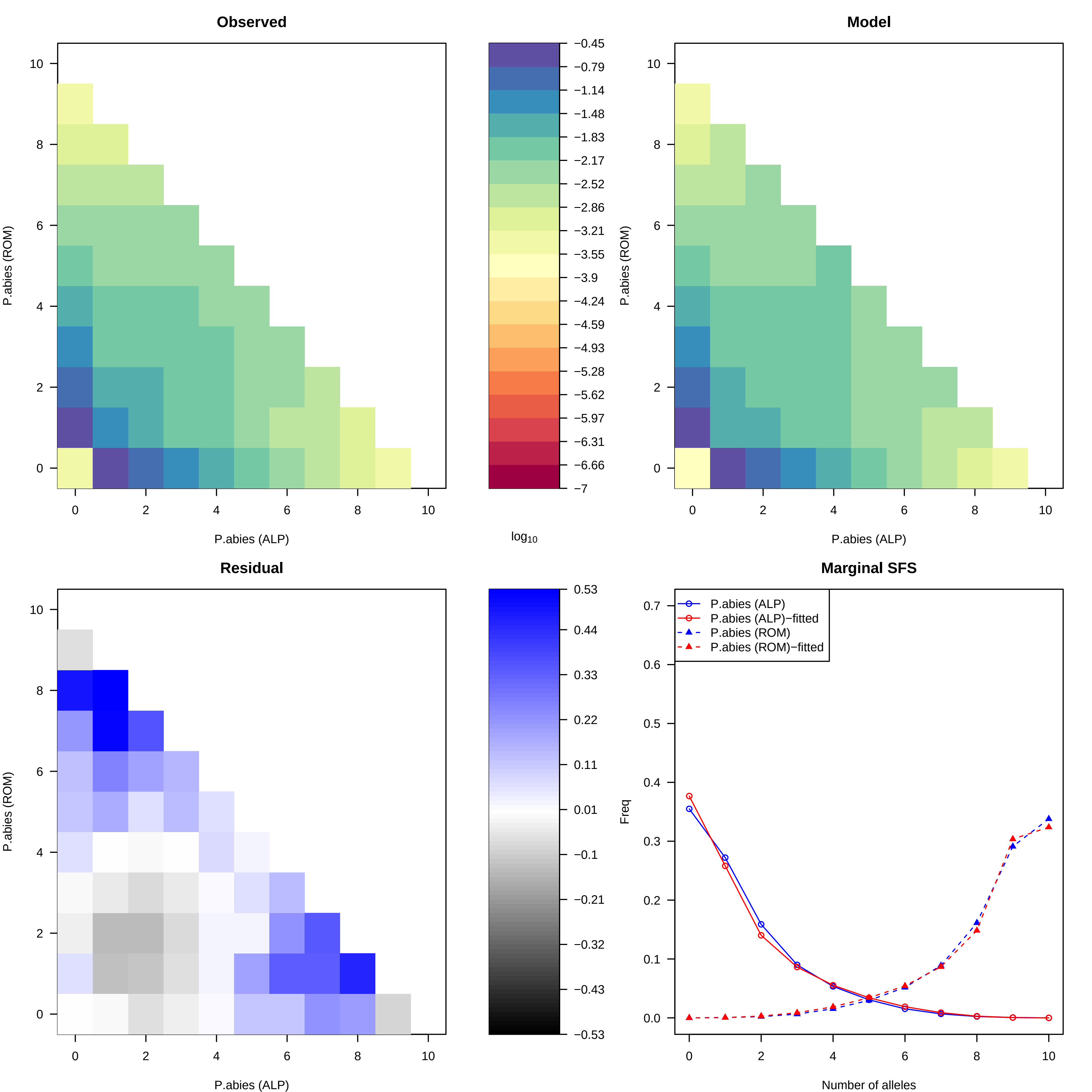
Observed
Model
Residual
Marginal SFS
P.abies (ROM)
P.abies (ALP)
log10
−0.45
−0.79
−1.14
−1.48
−1.83
−2.17
−2.52
−2.86
−3.21
−3.55
−3.9
−4.24
−4.59
−4.93
−5.28
−5.62
−5.97
−6.31
−6.66
−7
0.53
0.44
0.33
0.22
0.11
0.01
−0.1
−0.21
−0.32
−0.43
−0.53
P.abies (ALP)
P.abies (ALP)−fitted
P.abies (ROM)
P.abies (ROM)−fitted
Freq
Number of alleles
