## Supporting Information for "Genomic data provides new insights on the demographic history and the extent of recent material transfers in Norway spruce"

**Table S1: Demographic parameter estimates rescaled by a generation time of 50 years for *P. omorika (OMO)*, *P. obovata* (*OBO*), *P. abies* main domains and *P. abies – P. obovata* hybrids (*HYB*).**

| Parameters | Point estimation |
|---|---|
| $N_{OMORIKA}$ | 40 |
| $N_{OBOVATA}$ | 17,749 |
| $N_{HYBRID}$ | 200 |
| $N_{ALPINE}$ | 2,991 |
| $N_{CARPATHIAN}$ | 4,022 |
| $N_{FENNOSCANDIAN}$ | 3,770 |
| $T_{OMO\_OBO\_ABIES}$ | 22,875,400 |
| $T_{OBO\text{-}ABIES}$ | 17,600,050 |
| $T_{OBO\text{-}HYB}$ | 17,597,625 |
| $T_{FAC}$ | 15,274,375 |
| $T_{AC}$ | 15,272,700 |
| $TADM_{\_OBO\text{-}HYB}$ | 103,150 |
| $TADM_{\_OBO\text{-}ABIES}$ | 1,600 |
| $TBOT_{\_ABIES}$ | 12,850 |
| $TBOT_{\_OMORIKA}$ | 2,775 |

[a] Fennoscandian split from Alpine and Carpathian
[b] Alpine – Carpathian split

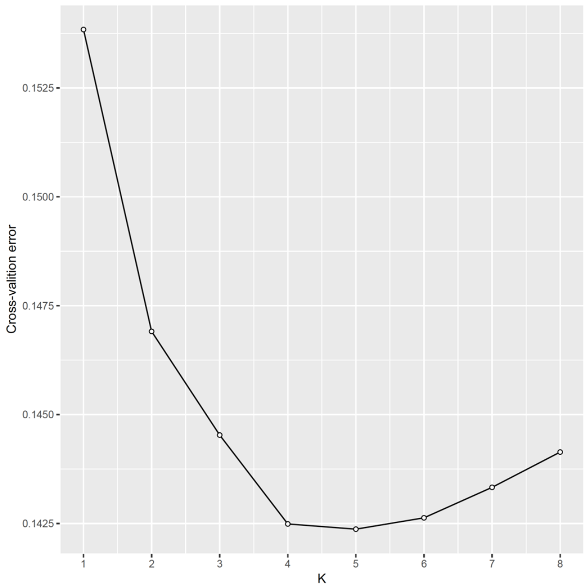


**Figure S1: Cross-validation error regarding number of cluster for unsupervised population clustering.**

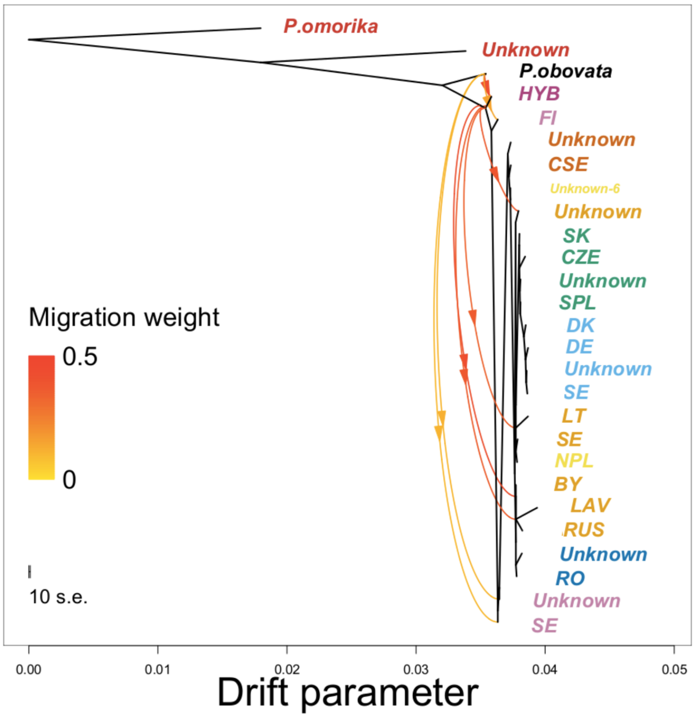


**Figure S2: TreeMix graph with 8 migration events.** Text colors showed the same genetic clusters in (Figure 2a and b). Russian-Baltic: Russia (RU), Belarus (BY), Estonia (EE), Latvia (LV), Lithuania (LT); Alpine: Germany (DE), Switzerland (CH), Denmark (DK), Sweden (SE); Central Europe: Slovakia (SK), Cze-republic (CZ), Southern Poland (SPL); Northern Poland (NPL); Romania (RO); Central Sweden (CSE); Fennoscandia: Finland (FI), Sweden (SE).

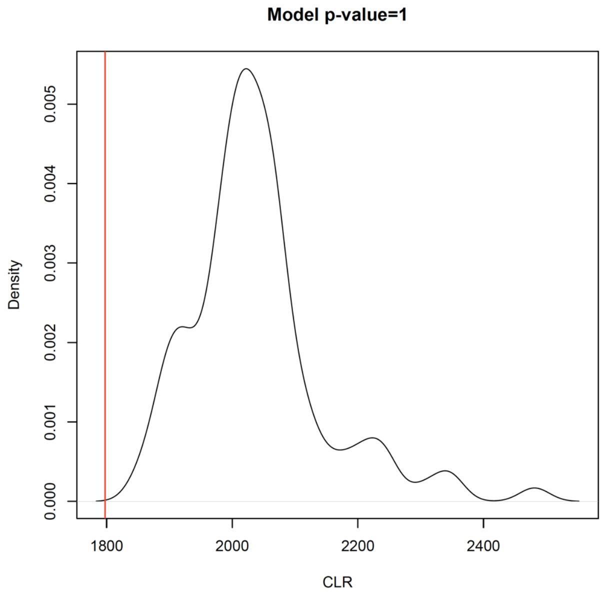


**Figure S3: Likelihood ratio G-statistics distribution**. The likelihood ratio G-statistics ($CLR = \log_{10}(CL_O/CL_E)$, where $CL_O$ and $CL_E$ are the observed and estimated maximum composite likelihood, respectively) was computed to evaluate model goodness-of-fit. A non-significant *p*-value of this test indicates that the observed SFS is well explained by the model. The red dotted line is the CLR of our divergence model.

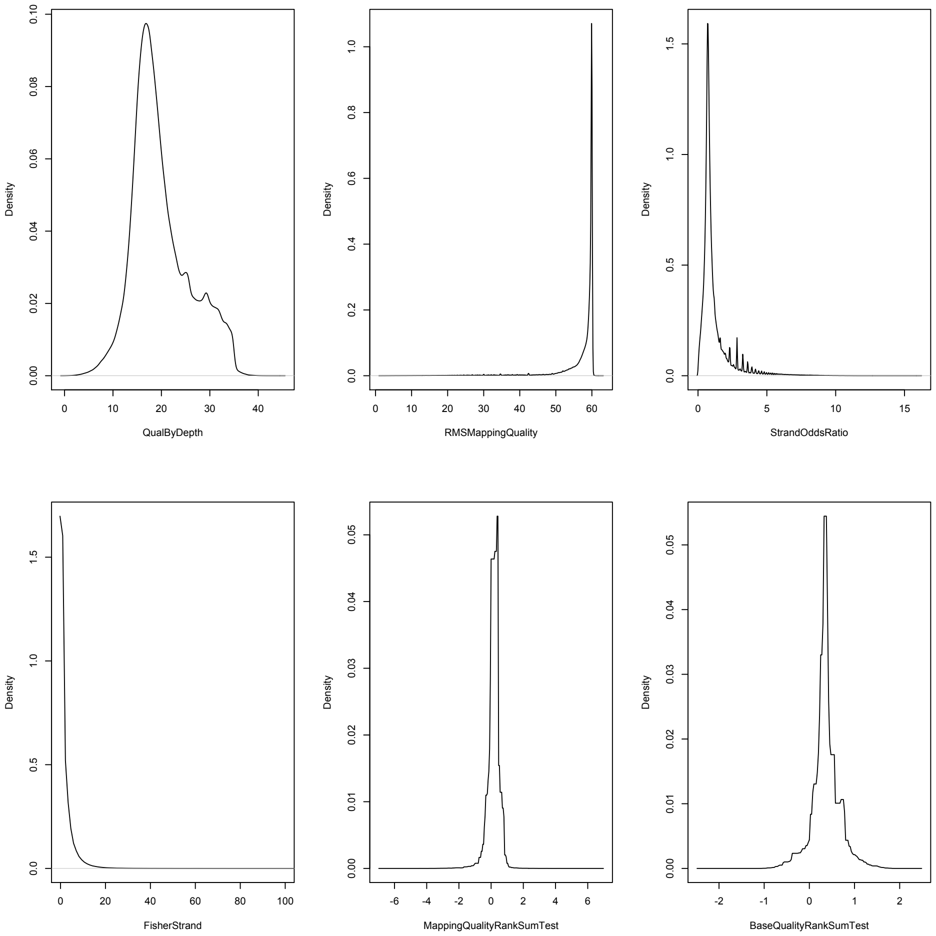


**Figure. S4 Variant quality scores reported for final SNP dataset after VQSR using GATK toolkit.** Density distributions of six quality scores (QD, MQ, SOR, FS, MQRankSum, and BaseQRankSum) were plotted to compare with generic recommendations (QD<2; MQ<40; FS>60; SOR > 3; MQRankSum < -12.5) for hard-filtering provided by Broad Institute.
